# Ligands binding to the cellular prion protein induce its protective proteolytic release with therapeutic potential in neurodegenerative proteinopathies

**DOI:** 10.1101/2021.04.19.440495

**Authors:** Luise Linsenmeier, Behnam Mohammadi, Mohsin Shafiq, Karl Frontzek, Julia Bär, Amulya N. Shrivastava, Markus Damme, Alexander Schwarz, Stefano Da Vela, Tania Massignan, Sebastian Jung, Angela Correia, Matthias Schmitz, Berta Puig, Simone Hornemann, Inga Zerr, Jörg Tatzelt, Emiliano Biasini, Paul Saftig, Michaela Schweizer, Dimitri Svergun, Ladan Amin, Federica Mazzola, Luca Varani, Simrika Thapa, Sabine Gilch, Hermann Schätzl, David A. Harris, Antoine Triller, Marina Mikhaylova, Adriano Aguzzi, Hermann C. Altmeppen, Markus Glatzel

**Author notes:** UCB Pharma SRL, Braine l’Alleud, Belgium. Authors contributed equally (Luise Linsenmeier, Behnam Mohammadi, Hermann C. Altmeppen). Corresponding authors &.

## Abstract

The cellular prion protein (PrP^C^) is a central player in neurodegenerative diseases caused by protein misfolding, such as prion diseases or Alzheimer’s disease (AD). Expression levels of this GPI-anchored glycoprotein, especially at the neuronal cell surface, critically correlate with various pathomechanistic aspects underlying these diseases, such as templated misfolding (in prion diseases) and neurotoxicity and, hence, with disease progression and severity. In stark contrast to cell-associated PrP^C^, soluble extracellular forms or fragments of PrP are linked with neuroprotective effects, which is likely due to their ability to interfere with neurotoxic disease-associated protein conformers in the interstitial fluid. Fittingly, the endogenous proteolytic release of PrP^C^ by the metalloprotease ADAM10 (‘shedding’) was characterized as a protective mechanism. Here, using a recently generated cleavage-site specific antibody, we shed new light on earlier studies by demonstrating that shed PrP (sPrP) negatively correlates with conformational conversion (in prion disease) and is markedly redistributed in murine brain in the presence of prion deposits or AD-associated amyloid plaques indicating a blocking and sequestrating activity. Importantly, we reveal that administration of certain PrP-directed antibodies and other ligands results in increased PrP shedding in cells and organotypic brain slice cultures. We also provide mechanistic and structural insight into this shedding-stimulating effect. In addition, we identified a striking exception to this, as one particular neuroprotective antibody, due to its special binding characteristics, did not cause increased shedding but rather strong surface clustering followed by fast endocytosis and degradation of PrP^C^. Both mechanisms may contribute to the beneficial action described for some PrP-directed antibodies/ligands and pave the way for new therapeutic strategies against devastating and currently incurable neurodegenerative diseases.

## Introduction

Neurodegenerative diseases, such as Alzheimer’s (AD) and Parkinson’s disease (PD) as well as less frequent prion diseases, not only share mechanisms of protein misfolding, protein aggregation and progressive spreading of pathology [1–3], but also involve common molecular players [4, 5]. One example is the cellular prion protein (PrP^C^), a highly conserved cell surface glycoprotein with high (yet not exclusive) expression in the nervous system [6].

Apart from its physiological functions, PrP^C^ plays a key role in prion diseases of humans (e.g. Creutzfeldt-Jakob disease (CJD)) and animals (e.g. chronic wasting disease in elk and deer; bovine spongiform encephalopathy in cattle). In these transmissible diseases, PrP^C^ misfolds into a pathogenic and partially proteinase K (PK)-resistant conformation (PrP^Sc^) [7, 8] due to either (i) a sporadic event, (ii) mutations in the coding *Prn-p* gene (causing genetic/familial disease forms) or (iii) contact with infectious “prions” (i.e., misfolded PrP species acting as “seeds” to template further PrP^C^ misfolding in acquired forms) [9]. More recently, GPI-anchored PrP^C^ has emerged as an important cell surface receptor for neurotoxic oligomers of β-sheet-rich peptides/proteins [6, 10–14] such as PrP^Sc^ itself, Aβ, tau and α-synuclein, which are all mediators of neuronal dysfunction found in neurodegenerative diseases such as prion diseases, AD, tauopathies, and PD, respectively [14, 15]. The plasma membrane is the primary site for the detrimental interactions of such extracellular toxic conformers with the disordered N-terminal part of signaling-competent PrP^C^ [16–18]. This binding causes synapto- and neurotoxic signaling (enabled by certain transmembrane proteins associating with PrP^C^ [19, 20]) and, in the case of PrP^Sc^ seeds, subsequent templated misfolding of native PrP^C^. In prion diseases, the survival time is inversely correlated with PrP^C^ expression levels [21, 22]. For these reasons, approaches to lower total or cell surface PrP^C^ levels are considered as promising therapeutic options with potential benefit also in the other abovementioned protein misfolding diseases [23–28].

Of note, surface levels of PrP^C^ are tightly regulated by various cellular mechanisms [29]. Among those is the proteolytic cleavage and extracellular release (shedding) by the metalloproteinase ADAM10 [30–32]. The latter is yet another example of a protein with relevance in different proteinopathies: Acting as the main ‘α-secretase’, ADAM10 is responsible for the non-amyloidogenic processing of the amyloid-3 precursor protein (APP), thus competing with the generation of toxic Aβ in the first place. As such, it has been proposed and investigated as a potential target in AD therapy [33–36]. Furthermore, by also lowering surface PrP^C^ levels, ADAM10 stimulation impairs the binding of Aβ to neurons and thus reduces toxicity [28]. In experimental prion diseases in mice, ADAM10 similarly confers protection as its expression correlates with survival time [37, 38]. Lastly, once being released from the surface, shed PrP (sPrP), which possesses all relevant binding sites, may interfere with PrP^Sc^ formation (in prion diseases) and block or neutralize various toxic conformers in the extracellular space [39]. Similar effects have already been described for recombinant or anchorless PrP versions, artificial PrP dimers, or the soluble N-terminal fragment (N1) resulting from the constitutive α-cleavage in the middle of PrP^C^ [11, 40–47], which may all be considered as a proxy for physiologically acting bona fide sPrP. In support of this, we here provide data obtained with murine disease models indicating that physiologically sPrP acts protective in neurodegenerative diseases by blocking and sequestering toxic oligomers.

When considering new treatment options against currently incurable diseases: why not utilizing a potentially protective process already provided by nature? While ADAM10 has been suggested as a therapeutic target in AD [33, 35], apparent problems arise from its rather broad expression pattern, the multitude of substrates in different tissues, and its involvement in various important physiological and pathological processes ranging from development and tissue homeostasis to intercellular communication and cancer [36, 48].

Therefore, directly manipulating this protease may cause significant side-effects, whereas a substrate-specific approach to stimulate the AD AM 10-mediated shedding of PrP^C^ would likely be superior.

Based on two earlier yet so far independent data-based concepts of (a) an increased ADAM10-mediated cleavage of some other ADAM10 substrates upon specific antibodybinding or dimerization [49–51], and (b) the protective effects of PrP^C^-directed antibodies in various models of Alzheimer’s and prion disease [52–66], we aimed at investigating how ligands binding to PrP^C^ would affect its supposed protective release by ADAM10. We show that a wide range of full-length (fl) IgG antibodies binding to central epitopes of PrP^C^ and some other PrP-directed ligands increase the ADAM10-mediated PrP-shedding. In contrast, a fl-IgG antibody targeting repetitive epitopes within the octarepeat region of PrP^C^ leads to strong PrP^C^ surface clustering and subsequent internalization and lysosomal degradation of the PrP^C^-antibody complex, whereas an identical derivative in its single-chain form increases shedding similar to abovementioned ligands. Moreover, we provide structural insight suggesting that shedding-stimulating effects of a PrP-directed antibody are enabled by a moderate conformational change in the relative positioning of the N- and C-terminal halves of PrP^C^.

Collectively, our data suggest that PrP^C^-to-ligand interactions play key roles in determining the fate of PrP^C^ regarding strong surface clustering followed by internalization and degradation on the one hand, or increased ADAM10-mediated shedding on the other hand. Both mechanisms may pave the way for novel therapeutic options against a wide range of dementias.

## Results

### Effects of ADAM10 and shed PrP in mouse models of neurodegenerative diseases

The role of ADAM10 in prion diseases has so far only been addressed by two studies in mice. Both studies, one using ADAM10-overexpressing mice [37] and one from our group employing a conditional, neuron-specific knockout model (A10 cKO; [38]), found a striking correlation between expression of the protease and survival times of prion-infected mice, thus pointing towards protective effects. In the latter study, we have also shown that PrP^Sc^ production was highest in A10 cKO mice, whereas only little PrP^Sc^ was detected in PrP-overexpressing *tga20* animals, even at a terminal stage of disease [38]. To directly assess sPrP and how this correlates with PrP^Sc^ amounts, we performed a biochemical comparison of brain homogenates (BH) of prion-infected A10 cKO mice and WT littermate controls (at 95 days post inoculation, dpi) as well as *tg*a20 mice (at 65 dpi, which in these mice corresponds to terminal disease). In agreement with our earlier study, highest PrP^Sc^ levels (upon sample digestion with PK) were found in A10 cKO, followed by moderate amounts in WT and lowest levels in *tg*a20 mice (Fig. 1A). Analysis of the respective non-digested samples –as expected for the genotypes– revealed that amounts of sPrP (assessed using our recently generated sPrP-specific sPrP^G228^ antibody [29]) were highest in *tg*a20 while hardly detectable in A10 cKO mice. This suggests an inverse correlation between this released PrP form and PrP^Sc^ production and may indicate that sPrP interferes with the conversion process. The seemingly contradictory fact that *tg*a20 mice, despite strongly elevated sPrP levels, still develop a rapidly progressive prion disease, is in line with recent studies [22] and supports the concept that the amount of membrane-associated PrP^C^ (which is increased in this model despite efficient shedding; [38]) rather than net PrP^Sc^ levels determine neurotoxicity. Next, we assessed localization of sPrP in terminally prion diseased *tg*a20 mice by immunohistochemistry. Despite exhibiting relatively low total amounts of PrP^Sc^ (Fig. 1A; [38]), deposits of PK-resistant PrP are mainly found in a layer between the hippocampus and corpus callosum (Fig. 1B, bottom picture). In contrast to the diffuse staining of sPrP in noninfected *tg*a20 brain described earlier [29] and shown for comparison in Fig. 2D, in prion disease sPrP is redistributed and clusters around those PrP^Sc^ aggregates (Fig. 1B), pointing towards close interaction between sPrP and PrP^Sc^ in respective deposits.

**Figure 1.**
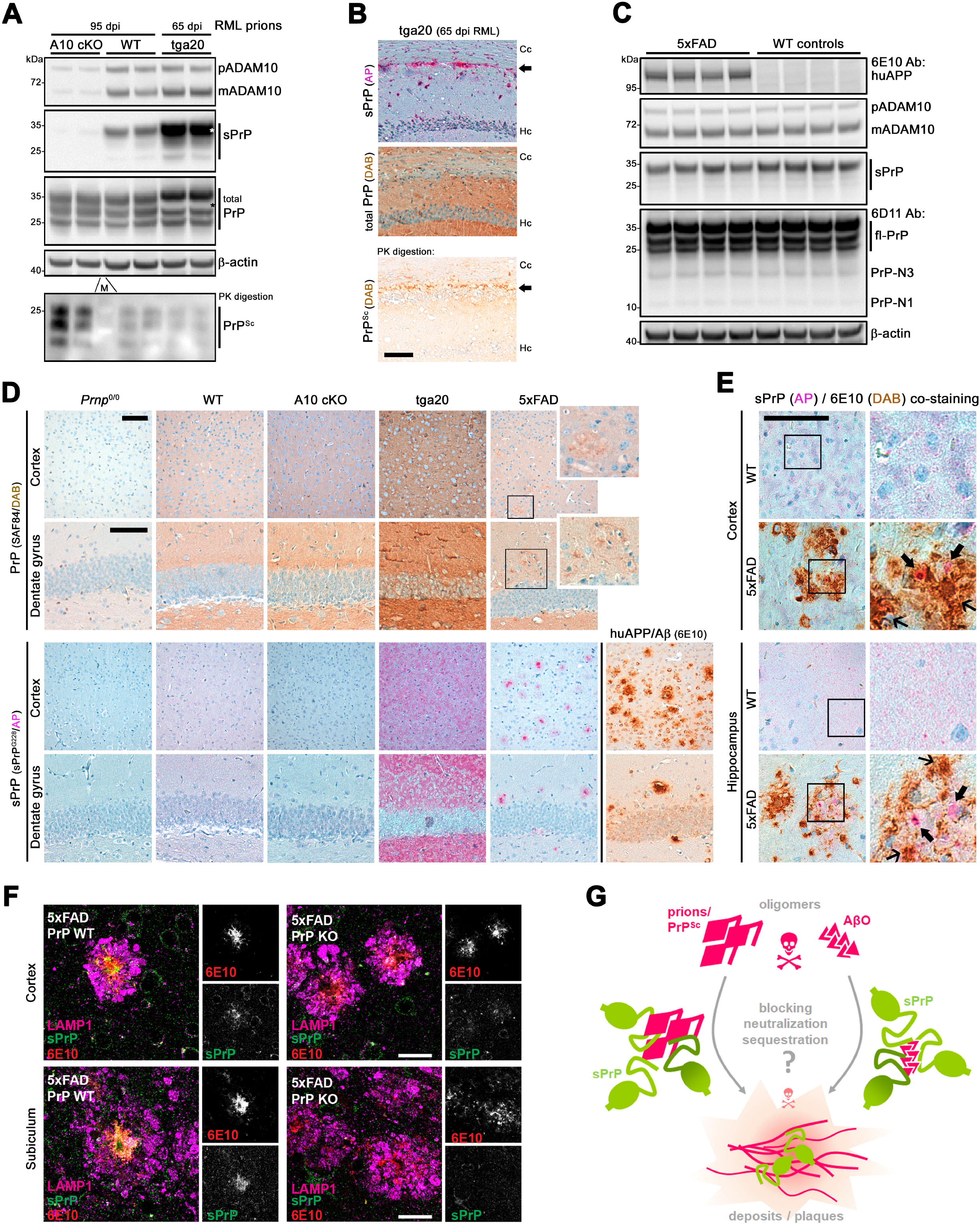
Shed PrP may interfere with toxic oligomers in neurodegenerative diseases. **a** Western blot analysis showing levels of premature (p) and active/mature (m) ADAM10, sPrP and total PrP in frontal brain homogenates of RML prion infected ADAM10 conditional knockout mice (A10 cKO) and wildtype (WT) controls (at 95 days post infection), and tga20 mice (at terminal disease; 65 dpi). Actin served as loading control. PK-resistant PrP^Sc^ in the same samples is shown in the lower blot. Asterisks mark position of the strong signal for diglycosylated sPrP in tga20 lanes causing a white (overexposed) area upon re-probing for total PrP. M = molecular weight marker lane. **b** Immunohistochemical (IHC) assessment of sPrP (pink signal due to detection with Alkaline Phosphatase (AP)), total PrP (brownish signal following detection with DAB) and PrP^Sc^ (upon PK treatment of section). Shown is the area between the CA region of the hippocampus (Hc) and the corpus callosum (Cc) of a prion infected tga20 mouse at a terminal state of disease. Arrows indicate the position of plaque-like PrP^Sc^ deposition (bottom picture) and a similarly clustered pattern of sPrP (top picture) contrasting with the regular diffuse pattern of sPrP observed in non-infected mice (shown in **d** for reference). **c** Biochemical assessment of ADAM10, sPrP and PrP (including N1 and N3 fragments resulting from endogenous α- and γ-cleavage, respectively) in the brains of 6-months old 5xFAD mice and WT littermate controls. Actin was detected as a loading control, human APP (using the 6E10 antibody) for genotype confirmation. **d** Representative IHC analysis showing PrP (brownish DAB signal; upper panel) and sPrP (pink AP signal; lower panel) in cortex and hippocampus (dentate gyrus) of a *Prnp*^0/0^ (negative control), WT, A10 cKO, tga20 and 5xFAD mouse. Plaque-like structures can be vaguely perceived in the pan-PrP staining (upper panel: boxes in 5xFAD samples and adjacent magnification). No sPrP signal is detected in *Prnp*^0/0^ and A10 cKO brain. Note that the diffuse pattern of sPrP (in WT and tga20) is converted to a clustered pattern in the 5xFAD brain similar to bona fide amyloid plaques detected in these mice (using the 6E10 antibody; right lower panel). **e** Co-staining of sPrP (AP) and amyloid plaques (DAB) in cortex and hippocampus of a WT and a 5xFAD mouse. Again, the diffuse pattern for sPrP (WT) changes to clustered signals in the 5xFAD sample. Bold arrows highlight pink signals of sPrP in rather loose Aβ deposits. Signal for sPrP is masked by the strong brownish signal in dense plaques (thin arrows). Scale bars in b, d, e are 100 μm. **f** Immunofluorescence (IF) staining of sPrP, Aβ plaques (6E10) and LAMP1 (as a marker for dystrophic neurites) on free-floating brain section showing cortex and subiculum of 5xFAD mice with (PrP-WT) and without PrP expression (PrP-KO). Note that sPrP colocalizes with Aβ plaques (in 5xFAD / PrP-WT), whereas only non-specific background is detected in the negative control (5xFAD / PrP-KO). Scale bar = 30 μm. **g** Scheme showing binding of sPrP (green) to toxic oligomers (pink). Sequestration of such extracellular oligomers into plaques (possibly supported by sPrP) may reduce neurotoxicity (indicated by sizes of the skull).

**Figure 2.**
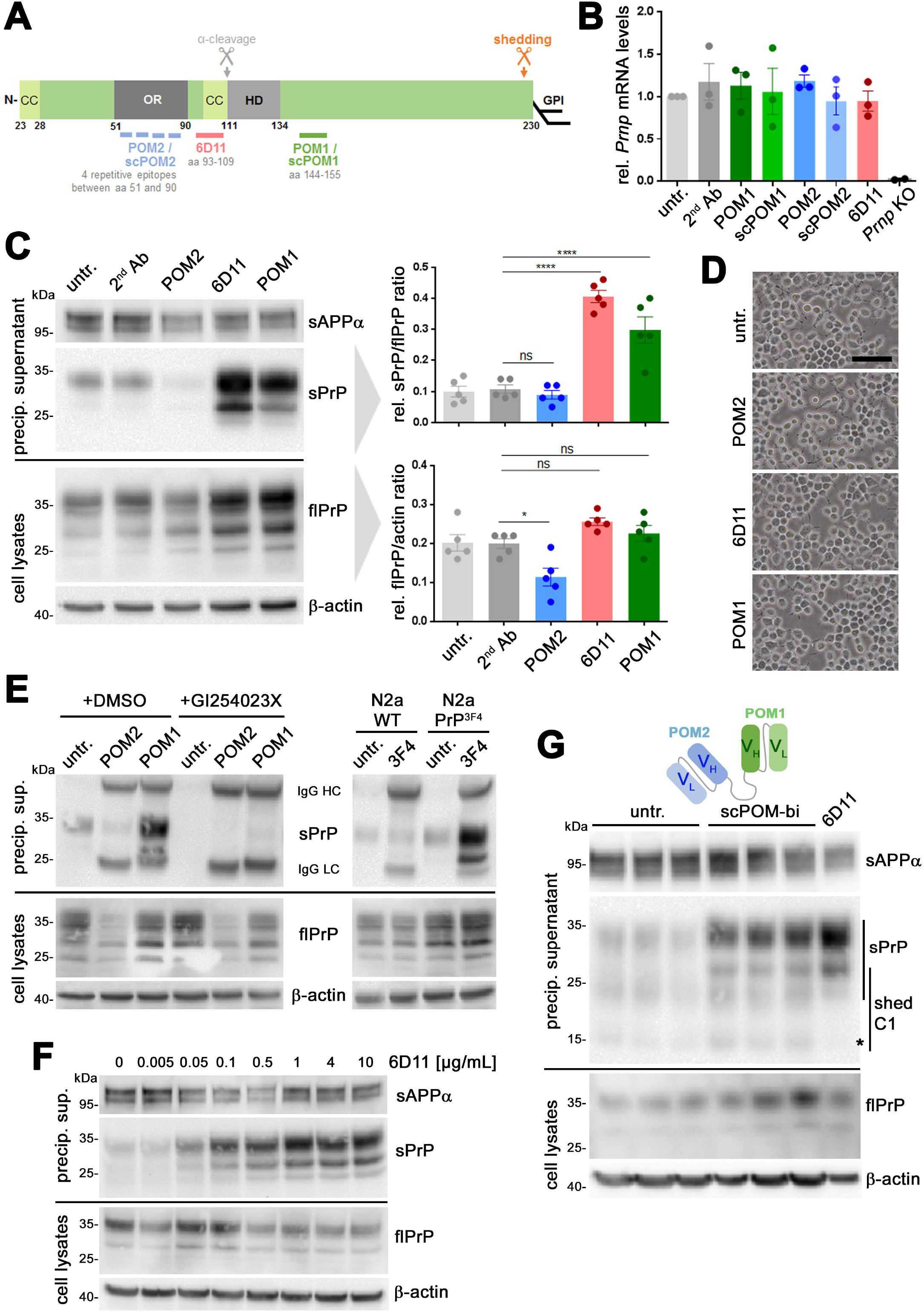
PrP-directed antibodies cause increased ADAM10-mediated shedding of PrP. **a** Linear representation of mature PrP showing important protein domains, position of the C-terminal GPI anchor, cleavage sites for shedding and α-cleavage, as well as epitopes of antibodies used in this study. CC: charged cluster; OR: octameric repeat region; HD: hydrophobic domain. **b** Quantification of *Prnp* mRNA levels in N2a cells either untreated (untr.) or treated for 16h with indicated antibodies or related single chain (sc) derivates. PrP-depleted N2a cells (*Prnp* KO) served as negative controls. n=3 independent experiments (n=2 for *Prnp* KO) each including 3 technical replicas. There were no statistically significant differences in the *Prnp* mRNA levels among different treatments and untreated controls. **c** Representative western blot analysis of full-length (fl) PrP in N2a cell lysates (lower panel) and sPrP (upper panel) in corresponding precipitated media supernatants after 16h of incubation with different PrP-directed IgGs. Actin (in lysates) and sAPPα (in media) served as loading controls. Quantifications of n=5 independent experiments are shown on the right. Average flPrP levels were found to be reduced (p≤0.05) only in POM2 treated cells compared to 2^nd^ Ab controls, however significant increase in the relative sPrP/flPrP levels was observed for 6D11- and POM1-treated cells compared to 2^nd^ Ab controls (p ≤ 0.0001). **d** Microscopy of untreated and treated N2a cells showing no alterations in density or overall morphology (scale bar = 100 μm). **e** Treatment of N2a cells with POM2 or POM1 antibody in the presence (+GI254023X) or absence (+DMSO; used as a diluent control) of a specific ADAM10 inhibitor (panel on the left). N2a WT cells or N2a cells stably expressing murine PrP with the human 3F4 sequence motif (N2a PrP^3F4^) treated or not with an antibody (3F4) targeting that sequence (panel on the right). Note that increased shedding is restricted to N2a PrP^3F4^ cells. **f** N2a cells treated with ascending concentration of the 6D11 antibody reveals a dose-dependency of the shedding-stimulating effect until reaching saturation at approximately 1 μg/mL. **g** Treatment of N2a cells with the bispecific immunotweezer (scPOM-bi) consisting of the complementarity-determining regions (V_H_ & V_L_) of both POM2 and POM1 antibody (scheme) causing increased shedding compared to untreated controls. Treatment with 6D11 was performed as positive control. Note that shedding of the PrP-C1 fragment is reduced upon 6D11 treatment (asterisk), which is likely due to 6D11 binding sterically hindering the PrP α-cleavage prior to shedding. Plotted data shows mean ± SEM for ‘n’ independent experiments. Statistical significance was estimated using ANOVA followed by Bonferroni’s multiple comparisons test with consideration of significance at \**p* < 0.05, \*\**p* < 0.005, \*\*\**p* < 0.001, \*\*\*\**p* < 0.0001.

In experimental models of Alzheimer’s diseases, released or recombinant forms of PrP exert protective effects by blocking neurotoxic Aβ oligomers (AβO) [11, 40–46, 67]. Stimulation of ADAM10 protects cells from AβO toxicity by reducing plasma membrane levels of PrP^C^ as receptor and toxicity transducer [28]. We aimed at assessing a potentially beneficial role of sPrP itself by blocking and sequestering toxic conformers in the extracellular space. Comparing levels of sPrP, total PrP and ADAM10 in brains of 6-month-old AD model mice (i.e., 5xFAD mice expressing human mutated APP among other manipulations) and WT littermate controls did not reveal differences in overall steady state levels of these proteins (Fig. 1C). In contrast to a previous report showing upregulation of the α-cleavage product PrP-N1 in human AD brain [45], we did not observe any similar effect in 5xFAD mice. However, upon histological assessment (Fig. 1D), the expected diffuse staining pattern for sPrP observed in WT and *tg*a20 mice shifted to a clustered distribution in 5xFAD brains strongly reminiscent of the bona fide amyloid plaques also detected in these mice (6E10-positive signals). As expected, *Prnp*^0/0^ and A10 cKO mice showed no sPrP signal. To exclude unspecific plaque-binding of our sPrP^G228^ antibody, we performed additional stainings comparing 5xFAD mice with matched PrP-depleted 5xFAD/*Prnp*^0/0^ mice (Suppl. Fig. 1). Moreover, when co-staining of sPrP and huAPP/Aβ was performed on 5xFAD and WT control sections, the diffuse signal for sPrP in the brain parenchyma observed in WT mice vanished at the cost of a more concentrated and plaque-associated pattern (Fig. 1E, thick arrows). Since presence of sPrP in dense plaques could not be addressed by IHC due to dominant brownish 6E10-positive DAB signal (Fig. 1E, slim arrows) and in order to gain insight at higher resolution, we additionally performed immunofluorescence (IF) stainings on free-floating sections (Fig. 1F). Indeed, sPrP clustered and co-localized within the cores of many (yet not all) 6E10-positive amyloid plaques surrounded by dystrophic neurites (indicated by the lysosomal marker LAMP1) (Fig. 1F). Again, besides weak unspecific background signal, no specific clustered pattern was observed with the sPrP^G228^ antibody in 5xFAD/*Prnp*^0/0^ mice used as negative controls (Fig. 1F).

Of note, it has repeatedly been described in human AD brain and respective mouse models that PrP^C^ (detected with pan-PrP^C^ antibodies) colocalizes with certain amyloid plaques and may even promote their formation [68–71]. Here we provide first evidence, that it is specifically sPrP (generated by ADAM10), which –as a diffusible factor in the extracellular space-is redistributed to the center of Aβ plaques in a murine AD model.

In sum, published data on protective effects of soluble released or recombinant PrP forms in prion diseases and AD, and published reports in addition to our novel findings shown here on the role of physiologically sPrP, support the view that sPrP blocks formation of (in the case of prion diseases) and sequesters/detoxifies (in prion diseases and other proteinopathies) harmful oligomeric protein conformers (scheme in Fig. 1G; [39]). This role of released PrP stands in clear contrast to the one of cell surface PrP^C^ acting as a toxicity mediator in these diseases [6, 11, 12, 14, 15].

### A substrate-specific approach to stimulate the ADAM10-mediated shedding of PrP

Earlier studies and our data presented above point to a protective role of PrP shedding in neurodegenerative diseases. Direct stimulation of ADAM10 is currently pursued in clinical AD trials albeit with the primary goal to stimulate the non-amyloidogenic processing of the amyloid precursor protein (APP) [35]. However, the multitude of ADAM10 substrates with (patho)physiological relevance throughout the body poses major challenges regarding potentially severe side-effects [36, 48].

For some ADAM10 substrates acting as cell surface receptors, binding of antibodies to their extracellular domains leads to their increased proteolytic release [49, 50]. Furthermore, as introduced earlier, PrP-directed antibodies show beneficial effects in different models of Alzheimer’s and prion diseases (reviewed in [72, 73]) and are even employed in the framework of a clinical trial [74]. These two seemingly ‘unrelated’ aspects prompted us to assess whether some selected PrP-directed antibodies (scheme in Fig. 2A) and ligands would stimulate its ADAM10-mediated shedding.

Treatment of murine neuroblastoma (N2a) cells with antibodies directed against central parts of PrP (6D11, recognising aa 93-109, and POM1, recognising aa 144-155 [75]) did not change overall PrP^C^ expression, as assessed by mRNA levels (Fig. 2B). However, levels of sPrP in media supernatants were significantly increased, whereas cell-associated PrP levels were unchanged (Fig. 2C). In contrast, treatment with an antibody directed against the flexible N-terminal part of PrP^C^ (POM2, recognizing repetitive epitopes between aa 51-90 [75]) led to a significant reduction in total PrP^C^ (Fig. 2C). Moreover, cell surface PrP^C^ levels, as assessed by a surface biotinylation assay, were increased upon POM1 and decreased after POM2 treatment (Suppl. Fig. 2).

Antibody treatment did not cause any obvious deleterious effects as judged by overall cell morphology (Fig. 2D) and an Annexin V apoptosis assay (Suppl. Fig. 3). To exclude cell line-specific effects, we also performed these experiments in another murine neuronal cell line (mHippo) and could essentially reproduce the observed effects regarding levels of sPrP, levels of cell-associated PrP (albeit with higher variation between experiments), and absence of overt toxicity (Suppl. Fig. 4).

As expected, co-treatment with the ADAM10 inhibitor GI254023X abolished the shedding-stimulating effect of POM1 confirming strict dependence of this process on this protease (Fig. 2E, left panel; [29]). This experiment also confirmed the reduction in total PrP^C^ upon treatment with POM2, which is independent of ADAM10 activity. We next treated cells with an antibody binding to an epitope in human PrP^C^ (3F4), which is absent in murine PrP^C^ (Fig. 2E, right panel). While no stimulating effect on shedding was observed in murine WT N2a cells, this antibody caused a strong increase of sPrP in N2a cells genetically modified to express 3F4-tagged PrP. Thus, the stimulated shedding observed here is executed by ADAM10 and specifically mediated by the binding of certain antibodies to PrP^C^.

We then addressed a potential dose-dependency of the shedding-stimulating effect using 6D11, which consistently caused the strongest stimulation among the PrP^C^-directed antibodies used in this study. Indeed, 6D11-mediated effects on PrP^C^ shedding are dosedependent with effects seen at concentrations as low as 0.005 μg/mL and reaching saturation at approximately 1 μg/mL (Fig. 2F).

Next, we tested a bispecific immunotweezer (scPOM-bi), which was recently shown to interfere with the formation of toxic prion species [76]. This chimeric antibody is composed of the complementarity-determining regions of POM1 and POM2. Being directed against both, the globular C-terminal and the flexible N-terminal half of PrP, this molecule may act intra- and intermolecularly, thus bridging the two dissimilar halves within one or between two PrP molecules, respectively. As shown in Fig. 2G, similar to POM1 and 6D11, treatment with scPOM-bi caused a clear increase in sPrP. Interestingly, shedding of the N-terminally truncated C1 fragment, which only displays one binding site for scPOM-bi, is only mildly increased when compared to non-truncated sPrP.

In sum, with the exception of POM2 (an anti-PrP^C^ antibody recognising repetitive N-terminal epitopes further investigated and discussed below), all PrP^C^-directed antibodies tested here caused a significant increase in the ADAM10-mediated shedding. As expected and demonstrated by unaffected levels of sAPPα (i.e., a cleavage product of APP also generated by ADAM10, Fig. 2C,F,G), this effect is substrate-specific.

### The PrP^C^-directed antibody 6D11 causes increased shedding in the absence of toxicity in organotypic brain slice cultures

After confirming the shedding-stimulating effect of some antibodies in two neuronal cell lines, we investigated whether this also holds true in a more complex biological system. We therefore tested the effects of the 6D11 antibody (as this IgG led to highest levels of sPrP in cell lines) in murine cerebellar organotypic slice cultures (COCS; [77]) derived from *tg*a20 mice. As expected, treatment with 6D11, but not with 3F4 antibody (an anti-PrP^C^ antibody not binding to murine PrP^C^ and therefore used as negative control), significantly increased levels of sPrP in the culture media surrounding the COCS (Fig. 3A, upper panel). No alterations in levels of full-length PrP^C^ were observed in the corresponding COCS homogenates (Fig. 3A, lower panel). Similar to the results obtained with N2a cells (Fig. 2C,F,G), the increase in PrP shedding in COCS upon treatment with 6D11 did not affect levels of sAPPα, thus confirming the substrate-specificity of this strategy (Fig. 3B).

**Figure 3.**
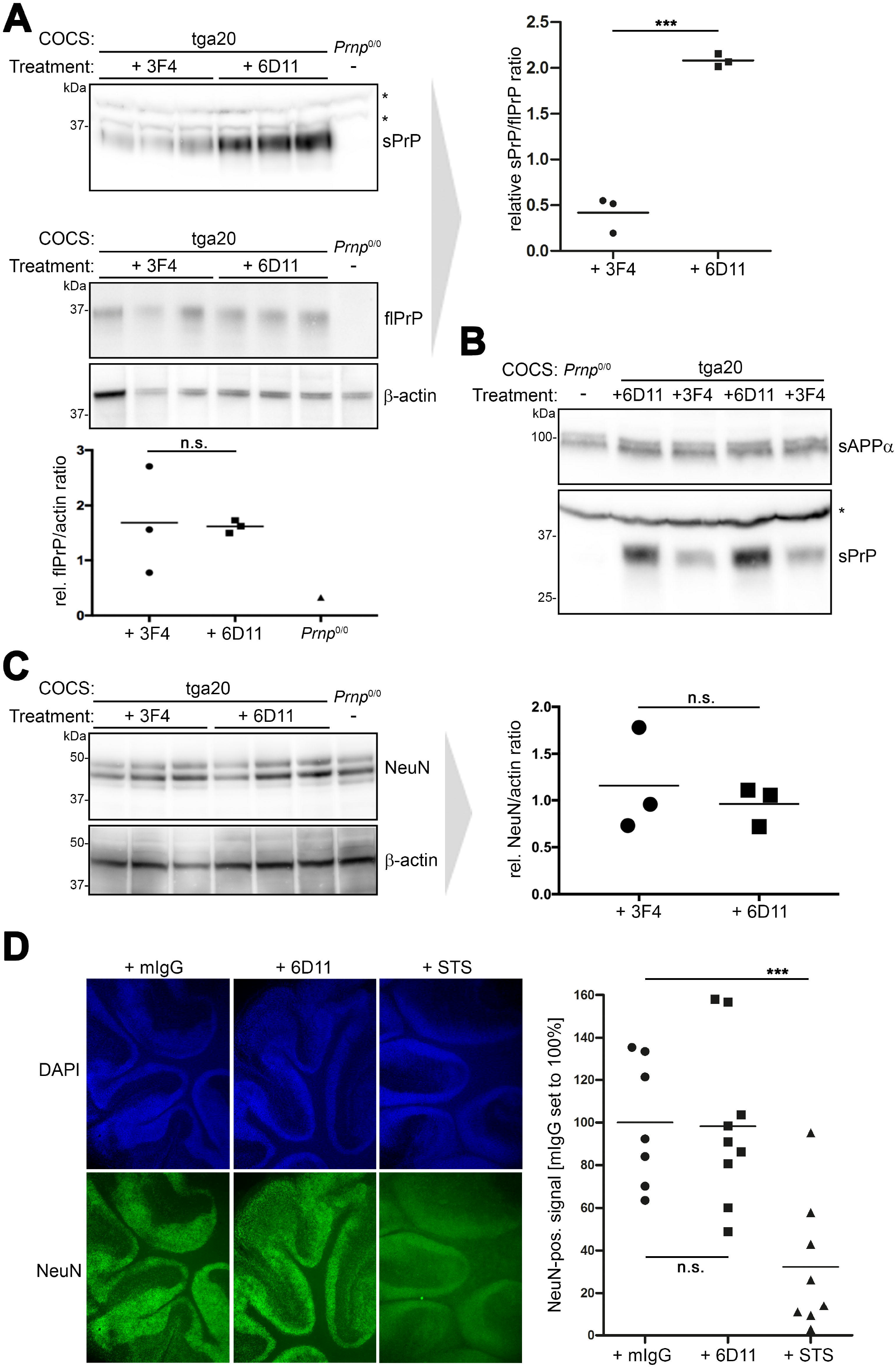
Stimulated shedding and lack of toxicity in antibody-treated murine organotypic brain slice cultures. **a** Cerebellar organotypic cultured slices (COCS) prepared from tga20 mice (or a *Prnp*^0/0^ mouse as negative control) and exposed to either 3F4 IgG (as negative control treatment) or 6D11 antibody. Western blot analysis showing levels of PrP in COCS homogenates (lower panel including quantification; actin was used as loading control) and sPrP in the culture media (upper panel with quantification on the right). **b** Biochemical assessment of sPrP and sAPPα in culture media after treatment as above. Asterisks in **a** and **b** indicate presence of unspecific bands (note the presence in *Prnp*^0/0^ samples) detected with the sPrP^G228^ antibody in COCS media. **c** Levels of the neuronal marker NeuN in above-mentioned COCS homogenates and densitometric quantification (actin used as reference). **d** Morphological analysis of antibody-treated COCS sections prepared from WT mice. Non-PrP-directed mouse antibodies (mIgG) were used as negative control, whereas staurosporine (STS) was used to induce toxicity and neuronal loss. DAPI staining (blue) reveals nuclei of all cells while NeuN staining indicates presence of neuronal nuclei. Representative sections are shown. Quantifications of the NeuN-positive signal are presented on the right (mIgG: n=7; 6D11: n=9; STS: n=8). Significance was assessed using unpaired two-tailed Student’s *t*-test (A,C) and one-way ANOVA with Dunnett’s post-hoc test (D).

We also addressed potential toxic side-effects caused by 6D11 antibody treatment. However, neither biochemical (Fig. 3C) nor morphological assessment (Fig. 3D) of the neuronal marker NeuN gave evidence for enhanced neuronal cell death in 6D11-treated COCS when compared to controls incubated with 3F4 antibody or non-specific murine IgGs, whereas this could readily be demonstrated in COCS treated with staurosporine as a positive control (Fig. 3D).

In summary, substrate-specific and non-toxic stimulation of PrP shedding can be achieved by treatment with anti-PrP antibodies, not only in neuronal cell lines but also in more complex biological settings such as COCS.

### Structural rearrangements in PrP caused by 6D11 binding

Antibody-mediated crosslinking induces proteolytic release of other ADAM10 substrates [49, 50]. Fittingly, artificial homodimerization of PrP^C^ was shown to result in increased proteolytic processing [78] and to protect cells from prion propagation [79]. Thus, given the elevated PrP^C^ surface levels found upon treatment with POM1 (Suppl. Fig. 2), we cannot exclude that dimerization and prolonged surface retention of PrP^C^ also play a role in induced shedding caused by POM1 and 6D11 IgGs. However, as demonstrated by the use of single-chain antibodies in the next paragraph, crosslinking is clearly not a prerequisite for stimulated shedding. Therefore, we aimed to gain further insight into shedding-stimulating mechanisms upon antibody binding at the structural level.

The 6D11 antibody has shown neuroprotective effects in various studies and turned out to be the most efficient shedding stimulator among the PrP^C^ ligands tested here. To assess the structure of the PrP-6D11 complex (Fig. 4B) and compare it to PrP alone (Fig. 4A), we employed state-of-the-art small-angle X-ray scattering (SAXS). This method allows for structural analysis in solution, and to obtain low-resolution three dimensional models. In our measurements, recPrP scattering was well fitted using a structured, mostly α-helical C-terminal domain and a flexible N-terminal tail partially flanking the globular domain in relatively close proximity (Fig. 4A, C, for more information see Suppl. Fig. 5 and Suppl. Table 1). In line with earlier SAXS data [80], this suggests presence of a shielding “cloud” formed by the N-terminal part surrounding the globular domain. Depending on the movement and position of the flexible tail at a given moment, this may regulate access of ADAM 10’s catalytic domain to the C-terminal cleavage site in PrP^C^ (framed scheme in Fig. 4C). In addition, the N-terminus was shown to interact with the plasma membrane [81–84], which may further block the membrane-proximate shedding event. Of note, for recPrP in complex with 6D11 IgG (here modelled bound to two recPrP molecules), we observed a much more extended N-terminal conformation of recPrP at a wider angle from the globular C-terminal domain (Fig. 4D). Thus, despite their own size, it appears possible that PrP-directed antibodies allow for better accessibility of ADAM10 by decreasing the sterical hindrance posed by the flexible tail of PrP.

**Figure 4.**
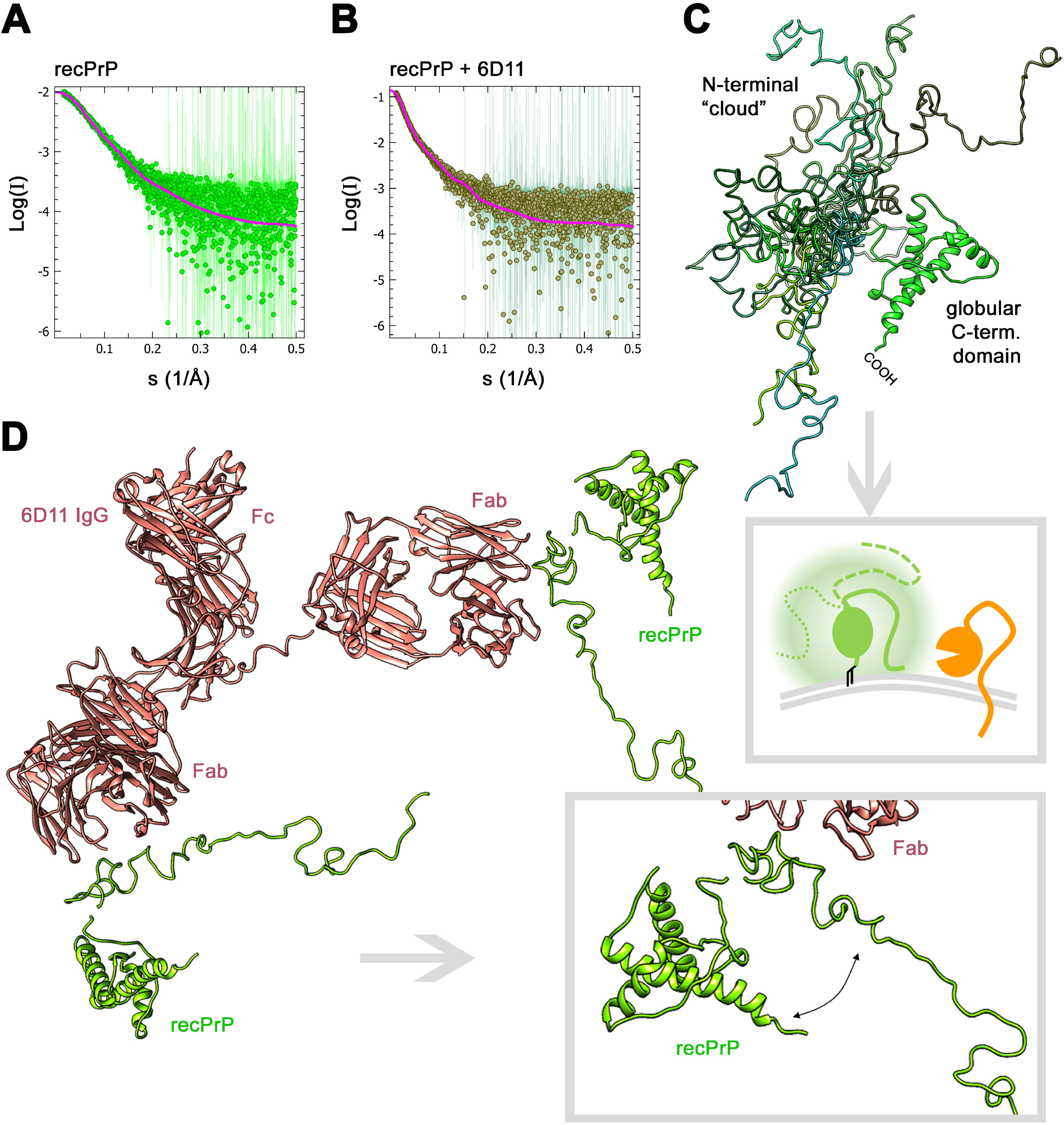
SAXS curves and modeling for recPrP (23–230) and the recPrP/6D11 antibody complex. Experimental SAXS profiles (dots) and fits (solid lines) for the best fitting model of recPrP; data source: SASBDB accession code: SASDHV9 (χ^2^=0.85; **a**) and the complex of two recPrP bound to 6D11 IgG (χ^2^=0.83; **b**). **c** Overlay of recPrP models resulting from SAXS measurements and showing multiple possible conformations of the flexible N-terminal tail (different shades of green) flanking the structured C-terminal domain. Framed scheme below outlines that movement of the flexible tail may create a “cloud” (shadowy corona) surrounding the globular domain and partially shielding PrP (green) from being shed by ADAM10 (orange), depending on the actual positioning and potential membrane interactions of the flexible tail (solid vs. intermitted lines). **d** Model of the recPrP (23-230) / 6D11 IgG complex. The magnified view (framed box) highlights an extended conformation of the flexible N-terminal region and an increased angle and distance to the C-terminal domain (as required to form a complex consistent with the SAXS data). Note that posttranslational modifications such as N-glycans and the GPI anchor are lacking in these analyses using recPrP.

### Stimulated shedding is also achieved by treatment with single-chain antibodies

For some ADAM10 substrates, such as CD44, antibody-mediated crosslinking causes increased cleavage [51]. Since, apart from POM2, all PrP^C^-directed IgGs tested here (POM1, 6D11; as well as 3F4 in respective cells expressing 3F4-tagged PrP^C^) and the bispecific immunotweezer (scPOM-bi) stimulated shedding, it was obvious to consider crosslinking of two PrP^C^ molecules as one underlying principle. In this scenario (and supported by the finding of increased PrP^C^ membrane levels caused by POM1 treatment mentioned above; Suppl. Fig. 2), dimerization would cause PrP^C^ to escape its usually high endocytosis rate [85] by stabilizing PrP^C^ and prolonging its presence at the cell surface, where ADAM10-mediated shedding is thought to occur. Accordingly, treatment with single chain antibodies, which are unable to crosslink two molecules, should not yield elevated sPrP levels.

To test for this, we treated N2a cells with single-chain versions of the two contrarily acting “ancestor” IgGs POM1 (scPOM1) and POM2 (scPOM2). Interestingly, both single-chain antibodies resulted in increased shedding compared to controls (Fig. 5A,B). Again, no toxic effects were observed based on overall cell morphology (Fig. 5C) and an Annexin V toxicity assay (Suppl. Fig. 3). To exclude stabilizing effects of these treatments on surface PrP levels, we again performed a surface biotinylation assay yet did not detect significant differences compared to the control treatment (Fig. 5D,E).

**Figure 5.**
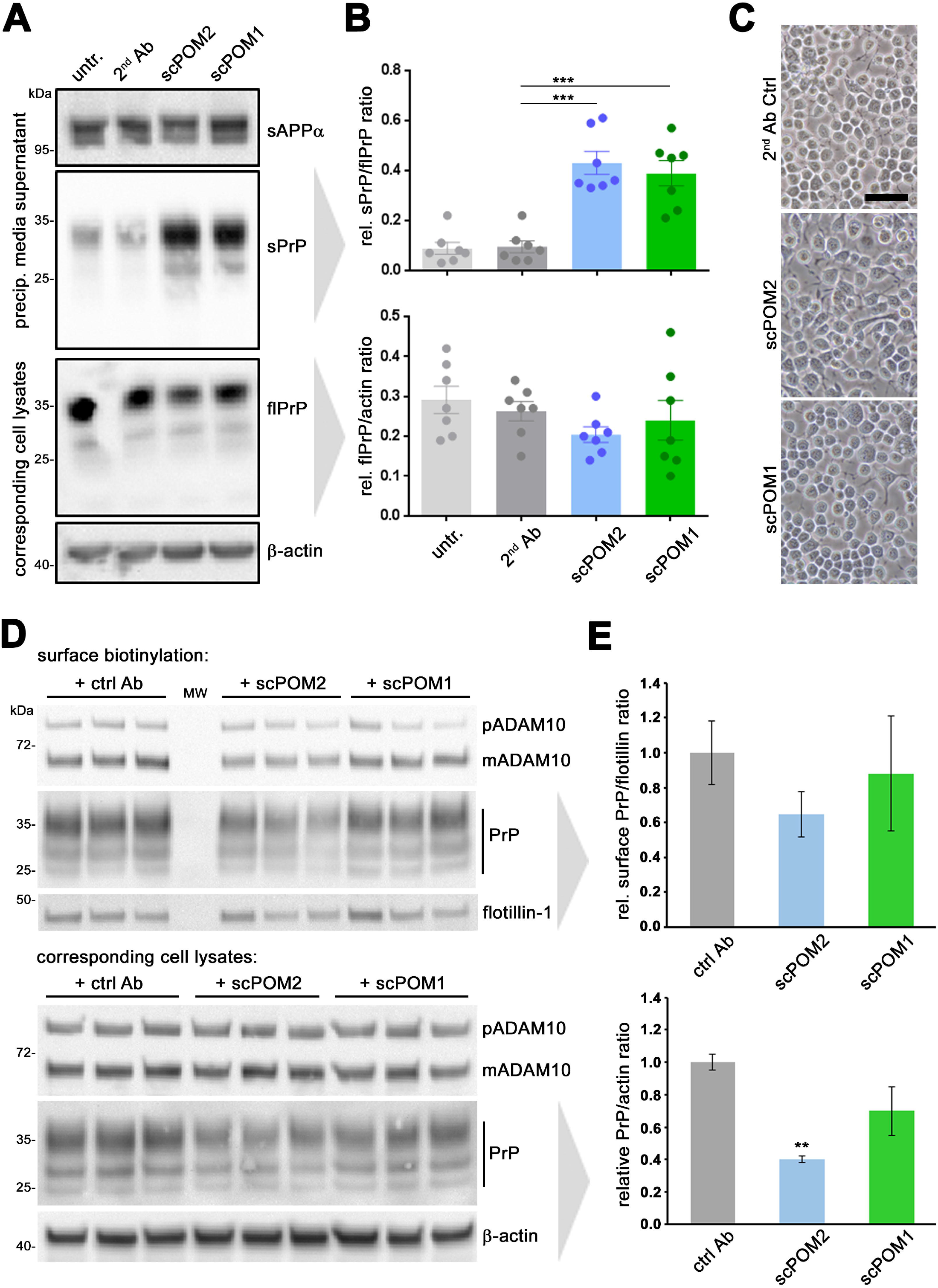
Single-chain antibodies induce shedding without causing PrP surface retention. **a** Representative western blot analysis showing levels of sPrP (and sAPPα as loading control) in precipitated media and flPrP (and actin as loading control) in respective lysates of N2a cells treated with single chain variable fragments of POM2 (scPOM2) and POM1 (scPOM1) antibodies. Untreated (untr.) and anti-mouse secondary antibody-treated (2^nd^ Ab) cells served as controls. **b** Densitometric quantification of sPrP (upper diagram) and cell-associated PrP levels (lower diagram). Plotted data shows mean ± SEM for n=7 independent repetitions. One-way ANOVA and Bonferroni’s multiple comparisons test was used to calculate significances. Relative sPrP/flPrP level was found to be increased in both scPOM-treated cells in comparison to that of 2^nd^ Ab controls (p ≤ 0.001). **c** Microscopic assessment of cell density and morphology (scale bar = 50 μm). **d** Cell surface biotinylation assay (upper panel) revealing membrane levels of ADAM10, PrP and flotillin. Total levels of ADAM10 and PrP in respective cell lysates are shown below. Actin served as loading control in lysates. Note the relative shift towards diglycosylated PrP and mature ADAM10 in biotinylated samples (compared to lysates) as these forms are thought to primarily locate at the cell surface. MW: lane used for molecular weight ladder. **e** Densitometric quantification of PrP levels presented in **d**. Plotted data shows mean ± SEM for n=3 technical replicas (shown in d). Significance was assessed using Student’s *t*-test (\*\**p* < 0.005).

These data strongly suggest that dimerization and an associated stabilization of PrP^C^ at the cell surface (as indeed detected for POM1 IgG; Suppl. Fig. 2) are -at least-not a prerequisite for the shedding-stimulating effect observed in this study. Our results also speak against an epitope-specificity of this effect. Instead, the combined data rather point towards a more general role of ligand-binding to PrP^C^ in stimulating its shedding. Moreover, it is intriguing that a single-chain version of POM2, despite addressing the very same epitopes, acts completely opposite to its IgG ‘ancestor’ with regard to the ADAM10-mediated shedding. This aspect will be further outlined below.

### Shedding is not stimulated upon treatment with four chemical compounds known to bind PrP^C^

The finding of stimulated shedding caused even by single-chain antibodies highlighted the possibility that various -and maybe even much smaller-ligands of PrP^C^ could likewise cause this effect. This would be particularly tempting considering potential therapeutic approaches and known difficulties associated with the use of antibodies in that regard (such as routes and doses of administration, costs, biostability, and passage through the blood-brain barrier). In an initial attempt, we therefore investigated four small chemical compounds shown to bind to different regions within PrP^C^ (highlighted in Suppl. Fig. 6A) and described to exert antiprion activity, at least *in vitro* (reviewed in [86]):

(i) GJP49, an anti-prion molecule identified through *in silico* analyses aimed at directly identifying pharmacological chaperones for PrP^C^ [87]; (ii) Chlorpromazine, an anti-psychotic drug originally claimed to inhibit prion propagation by directly binding to PrP^C^ [88] but more recently shown to rather promote re-localization of PrP^C^ from the cell surface [89]; (iii) The porphyrin Fe(III)-TMPyP, perhaps the only extensively validated PrP^C^ ligand, reported to inhibit prion propagation in a strain-independent fashion [90, 91]; and lastly (ii) Quinacrine, another tricyclic acridine derivative traditionally used as an anti-malaria drug and then identified as an anti-prion compound capable of directly binding to PrP^C^ [92, 93].

These candidates were tested using ascending concentrations (0.1, 0.3, 1, and 3μM) for treatment (24h) in an established screening system using HEK293 cells stably expressing murine PrP^C^ (Suppl. Fig. 6B; [94]). None of these compounds significantly altered levels of cell-associated PrP^C^. Likewise, no relevant changes for sPrP were detected with the surprising exception of Quinacrine, which rather caused a reduction in shedding. This could be due to its expected binding region at the C-terminus of PrP^C^ (Suppl. Fig. 6A) overlapping with the cleavage site and, thus, potential hindrance of the protease. However, since we were rather interested in ligands increasing the shedding, we did not further investigate this aspect. Although we are continuing to screen for small compounds that stimulate shedding, the findings presented here may indicate that ligands of PrP^C^ are required to exceed a critical size or fit certain sterical characteristics in order to stimulate its proteolytic shedding.

### The PrP^C^-reducing effect of POM2 IgG is linked to its special binding characteristics

Among the antibodies tested here, POM2 IgG (yet not its single-chain variant) represents an exception as it did not cause increased shedding but rather reduced levels of cell surface and total PrP^C^. This is interesting given that lowering (cell-associated) PrP^C^ amounts is considered one of the most promising strategies for prion disease therapy [22, 27, 95, 96]. Moreover, POM2 was shown to be neuroprotective as it impairs detrimental interactions of the flexible N-terminus with the plasma membrane caused by prions [81, 83, 84].

To further investigate the decrease in PrP^C^ caused by treatment with this antibody, we treated N2a cells with POM2 and assayed total PrP^C^ levels by western blot over time. PrP^C^ amounts were reduced as early as 15 min following POM2 treatment, reaching a stable plateau at 60 min and later (Fig. 6A). Since a reduction in cell-associated PrP^C^ with no parallel increase in media supernatants (Fig. 2C,E) indicates enhanced cellular degradation, we performed antibody treatments in the presence or absence of Bafilomycin, an antibiotic targeting the vacuolar H^+^-ATPase, thus impairing lysosomal acidification and functioning (Fig. 6B). This co-treatment had no drastic effect on cells treated with unspecific control antibody (Ctrl) or 6D11 (apart from a moderate increase and slightly altered banding pattern of particularly diglycosylated PrP suggesting effects of bafilomycin treatment on PrP glycosylation) yet was able to restore PrP^C^ levels in cells treated with POM2. This indicated that POM2 treatment leads to uptake and lysosomal degradation of PrP^C^ rather than induced shedding.

**Figure 6.**
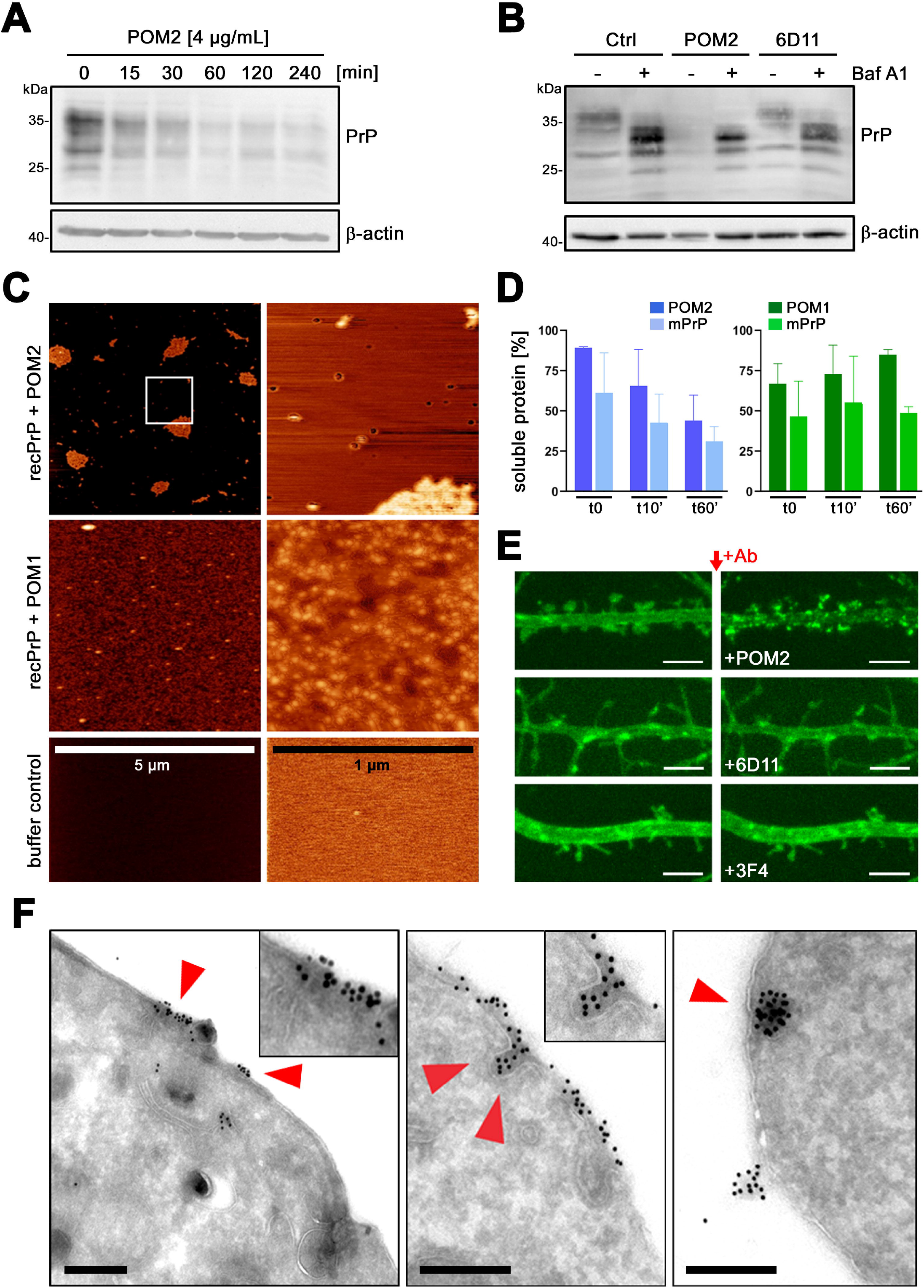
POM2 IgG treatment results in strong (surface) clustering, uptake and degradation of PrP. **a** Time-course experiment showing cell-associated PrP levels in N2a cells lysed at different time-points after treatment with POM2. **b** Western blot showing cellular PrP levels upon treatment with POM2, 6D11 or control antibody in the presence (+) or absence (-) of the lysosomal inhibitor Bafilomycin (Baf A1). Actin served as loading control in **a** and **b**. **c** Atomic force microscopy of recombinant PrP (recPrP) incubated with POM2 or POM1 antibody in overview (left column; scale is indicated) and fivefold further magnification (right column). Lower panel shows mica surface treated with protein-free buffer only. **d** Quantification of a solubility assay of a mixture of mouse recPrP (mPrP) with either POM2 (blue graphs) or POM1 (green graphs). Recovery of respective antibodies or PrP alone in solution immediately (t0), 10 min (t10’) or 60 min (t60’) after mixture was set to 100%. **e** Microscopic picture details showing dendrites of rat hippocampal neurons expressing GFP-tagged PrP (green) taken shortly before (left column) and after (right column) treatment with POM2, 6D11 or 3F4 antibody (for a complete overview refer to Suppl. Fig. 8 and Suppl. Movie 1). Scale bars represent 4 μm. **f** Representative electron microscopy pictures of N2a cells after 5 min (left) or 30 min (middle and right) of treatment with POM2 antibody showing clustering and uptake of PrP-directed immunogold particles. Scale bars represent 250 nm.

We reasoned that the exceptional behaviour of POM2 must result from its specific binding characteristics. In contrast to all other antibodies assessed here, POM2 can bind to four repetitive epitopes all located in the structurally disordered N-terminal part. Hence, POM2 IgGs (but not scPOM2) with their double valency and four epitopes, supported by the extreme structural flexibility in this region of PrP^C^, might be able to cluster large complexes of several PrP^C^ (and antibody) molecules. To test for this *in vitro,* we performed atomic force microscopy (AFM) of recombinant PrP incubated with either POM2 or POM1 antibodies (Fig. 6C). In contrast to the latter, which -at best-is able crosslink two PrP molecules and causes a dotty staining on the mica surface, POM2 treatment resulted in the formation of large clusters (on cost of the dotty pattern observed for POM1/recPrP).

We next confirmed these findings by a solubility assay (Fig. 6D), were recovery of both components in the soluble fraction of a mixture of either POM2 or POM1 with recombinant murine PrP was assessed over time (normalized against respective POM antibody or mPrP alone set to 100%). A clear trend of a progressive decrease of both components was observed for POM2, yet not for POM1, thus further supporting the abovementioned AFM findings of strong aggregation of POM2/recPrP complexes.

To investigate whether comparable clusters are also formed at the surface of cells upon POM2 treatment, we performed live microscopy of primary rat neurons transfected with GFP-tagged PrP^C^. As shown in the representative snapshots in Fig. 6E and the respective overview presented in Suppl. Fig. 7A and Suppl. Movie 1, POM2 treatment indeed caused a fast and strong surface clustering of PrP^C^, which was not observed with 6D11 or 3F4 antibodies used as controls. To exclude that this clustering by POM2 at the neuronal surface was only due to transgenic overexpression (GFP-PrP), we also performed staining of endogenous PrP^C^ after treatment with POM2 and found a comparable pattern of clusters (Suppl. Fig. 7B).

To address consequences of this clustering at a subcellular resolution, we also employed electron microscopy analysis upon immunogold labeling. As early as 5 min after incubation of N2a cells with POM2, we found large immunogold-positive clusters of PrP^C^ at the cell surface and many instances revealing subsequent endocytosis of these complexes (as represented in Fig. 6F), thus supporting POM2-stimulated uptake and degradation of PrP^C^ (Fig. 6A,B). In another set of experiments, this time directly comparing POM2-with POM1-treated cells, we confirmed cluster formation for POM2, yet could not find evidence for this in cells incubated with POM1 antibody (Suppl. Fig. 8).

Membrane proteins are highly dynamic due to lateral diffusion on the plasma membrane [97]. Clustering on the plasma-membrane is either a 2D (complex formation between laterally diffusing membrane proteins) or 3D event (complex formation between membrane protein and scaffolds). Using SPT-QD (Single Particle Tracking using Quantum Dots) in primary neurons, we monitored the events leading to PrP^C^ clustering by POM2 IgG. SPT-QD is a powerful method that allows for the measurement of lateral diffusion and protein-protein interaction at a single molecule resolution. Besides other POM-IgGs used as controls, we here also included POM11, which shares two important features with POM2: (i) having more than just one epitope, which are (ii) located within PrP’s flexible tail (FT). First, we measured the diffusion coefficient of endogenous PrP^C^ using 5 different POM antibodies (POM-x: POM1/19 (single epitopes within the C-terminal half / globular domain (GD)); POM2/POM11 (FT-binders with ≥2 epitopes as described above) or POM3: epitope in the central part / hydrophobic core (HC)) without prior exposure to the same POM-x antibodies (Fig. 7A). Synapses were labelled using FM4-64 dye and single molecule trajectories were obtained and analysed both in and out of synapses. Compared to the extra-synaptic sites, PrP^C^ diffusion was generally slower at the synapses, which is likely due to the crowded environment. Importantly, similar diffusion coefficient values of PrP^C^ were obtained with all POM antibodies (Fig. 7B, each data point represents an average value measured for hundreds of QDs; also refer to Suppl. Table 2). Thus, without prior exposure to high POM-x concentrations, PrP^C^ mobility remains largely unaltered. Next, neurons were exposed to POM-x antibodies (1 μg for 1h) prior to PrP^C^ diffusion measurement (Fig. 7C). Intriguingly, in neurons pre-exposed to FT-directed IgGs with two (POM11) or four (POM2) repetitive epitopes, diffusion of PrP^C^ showed a 20-60% slow-down (most pronounced for POM2). Contrary, pre-exposure to the other IgGs had no (POM3/POM19) or only moderate effects (<10% for POM1) on PrP^C^ diffusion. Each data point represents the difference (mean +/− SEM) between unexposed control and POM-x-exposed condition for a given experiment. The absolute diffusion coefficient values for all QDs are provided in Suppl. Table 2. In sum, the strong impact of POM2 IgG (and –albeit to lesser extent– the closely related POM11 binding to overlapping epitopes) on PrP mobility supports the idea that these IgGs drive freely diffusing PrP^C^ to form larger complexes, resulting in cluster formation.

**Figure 7.**
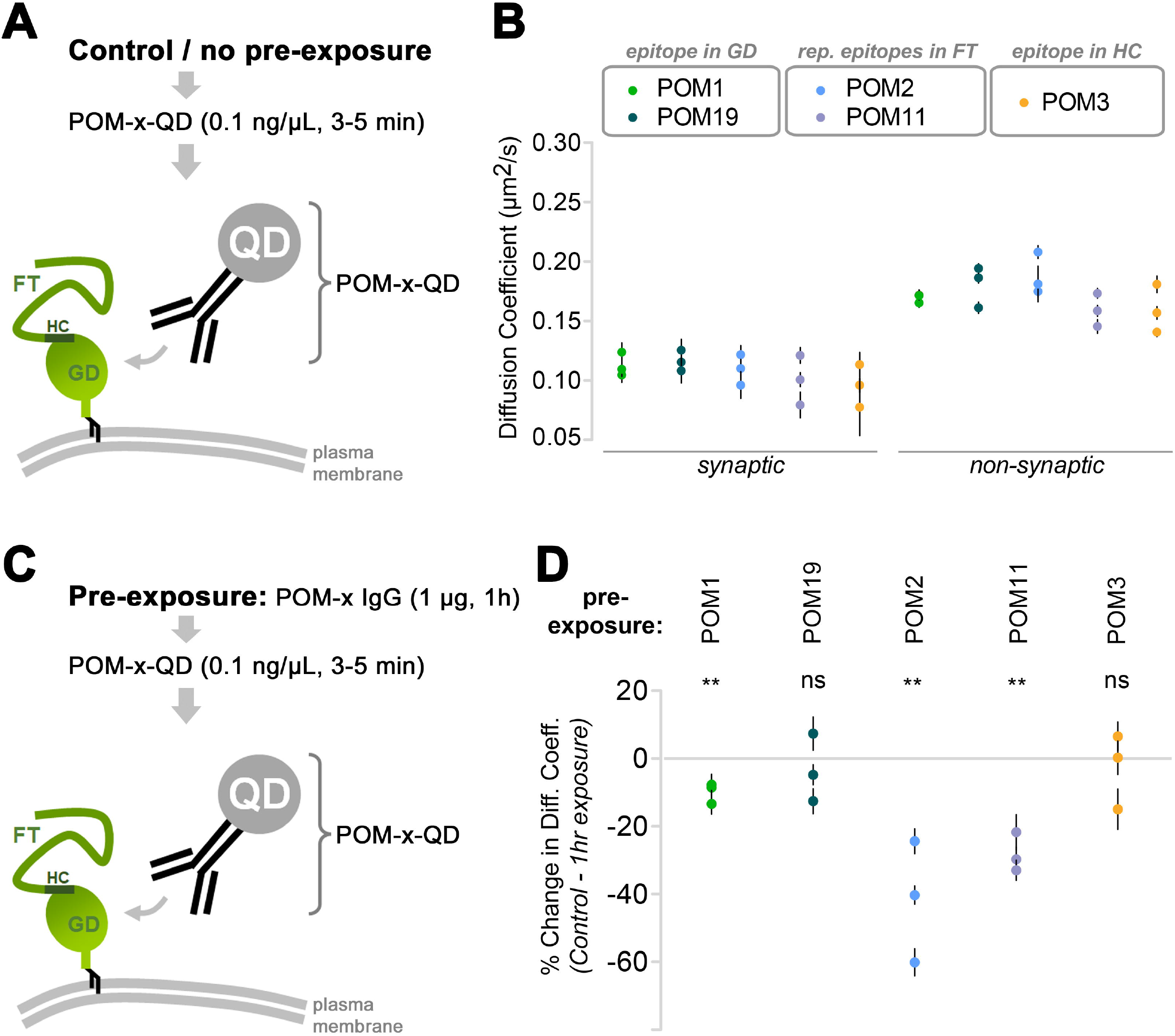
Slow-down in the lateral diffusion of endogenous PrP^C^ by IgGs directed against repetitive epitopes within the flexible tail. **a** Single particle tracking using Quantum Dots (SPT-QD) to quantify the diffusion coefficient of endogenous PrP^C^ and using QD-pre-coupled to anti-PrP^C^ antibodies (POM-x-QD). **b** Under control conditions (no preexposure to high concentration of antibodies), similar diffusion coefficient values of PrP^C^ was obtained using various antibodies (globular domain (GD)-directed IgGs: POM1 and 19; flexible tail (FT)-directed IgGs: POM2 and 11; or hydrophobic core (HC)-directed: POM3). Synapses were identified using FM4-64 labeling. Plotted data shows mean ± SEM values for 3 independent experiments and One-way ANOVA test was performed with no significant difference. **c** SPT-QD of PrP^C^ using POM-x-QD antibody following pre-exposure (1h) to high concentration (1 μg) of POM-x antibodies. **d** Only pre-exposure to FT-directed IgGs but not to the others greatly (>20%) reduced the diffusion coefficient of PrP^C^. Plotted data shows mean ± SEM for 3 independent experiments. Paired *t*-test was performed to compare the difference from control condition (no pre-exposure, **b**). Data for all QDs analysed is shown in Suppl. Table 2.

## Discussion

A soluble form of PrP, most likely representing sPrP, has been described decades ago [30]. However, convenient and reliable discrimination between sPrP and the usually much higher (and thus masking) amounts of membrane-associated fl-PrP^C^ in biological samples has only been achieved recently, fueled by the generation of antibodies specifically recognizing sPrP [29, 98–100]. Thus, knowledge on potential functions of sPrP is mainly based on data obtained using recombinant PrP (recPrP). Although recPrP lacks glycosylation and may thus differ structurally from both, PrP^C^ and physiologically released sPrP, data obtained using recPrP imply physiological functions in synapse formation [101], neurite outgrowth guidance [102], differentiation, immune signaling and intercellular communication [103–105]. Importantly, one of the few molecularly well-defined physiological roles of PrP^C^ (i.e., in regulating myelin maintenance in the peripheral nervous system) might in fact be executed by sPrP (and/or PrP’s released N1 fragment) [106].

Regarding a role of sPrP in prion diseases, data show that released forms of PrP, secreted PrP dimers, but also recPrP confer neuroprotection mainly by blocking the buildup of neurotoxic protein conformers [40, 107–109]. Indeed, we and others have already shown that levels of the PrP sheddase ADAM10 correlate with incubation times in prion infected mice, and provided indirect evidence for an inverse correlation between ADAM10 expression (and, thus, supposed levels of sPrP) on the one hand, and neurotoxic PrP^Sc^ formation on the other hand [37, 38]. Using our sPrP-specific antibody, we here provide more direct evidence for this. Interestingly, we found sPrP to co-localize with PrP^Sc^ in respective deposits in prion diseased mice. This may indicate that sPrP binds to neurotoxic PrP^Sc^ in the extracellular space and helps to sequester those conformers into deposits (Fig. 8A), and could also explain the fact that even mild transgenic overexpression of ADAM10 leads to a relevant decrease in PrP^Sc^ production in the brain [37].

**Figure 8.**
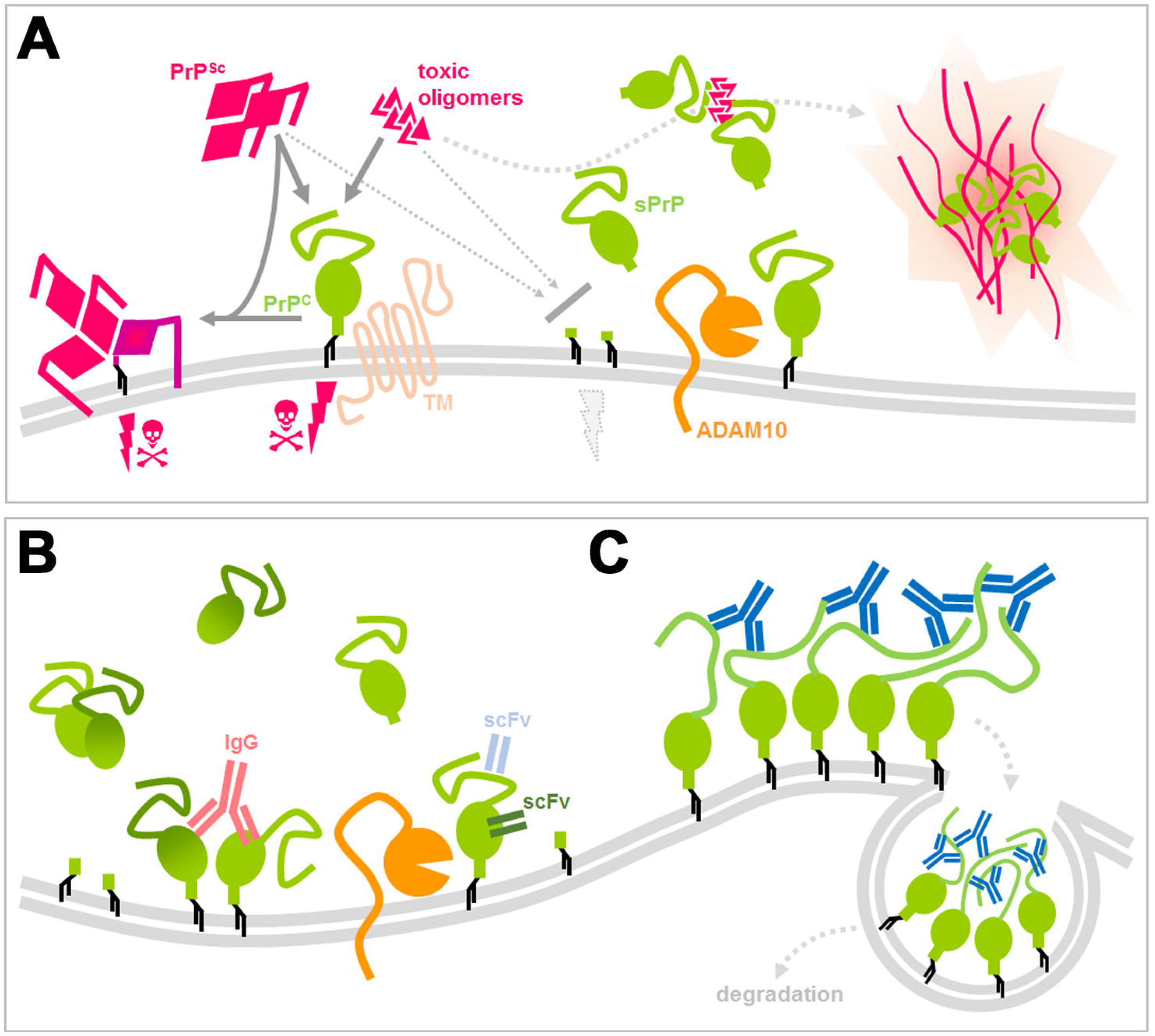
Scheme summarizing potential protective roles of sPrP and effects of PrP-directed ligands. **a** PrP (green) expressed at the cell surface is a central player in neurodegenerative diseases as it serves as a substrate for prion conversion and PrP^Sc^ (pink) production in prion diseases and (in complex with certain transmembrane proteins) acts as a receptor for toxic protein conformers (pink), such as Aβ in AD, initiating toxic signalling (pink thunderbolts and skulls). As supported by several published reports, mechanisms that lower PrP levels at the plasma membrane, such as the endogenous shedding mediated by ADAM10 (orange), are considered neuroprotective. In addition, several studies have shown that released forms or fragments of PrP interfere with toxic proteins in the extracellular space. Our data suggest that sPrP inversely correlates with PrP^Sc^ formation and colocalizes with deposits of PrP^Sc^ and Aβ, indicating a blocking and possible sequestrating activity of sPrP towards harmful conformers. As a consequence, stimulated shedding may represent a promising therapeutic option. **b** Regarding the latter, we here provide evidence that several PrP-directed ligands binding to different epitopes cause an increased ADAM10-mediated shedding. While crosslinking may be involved (as in the case of IgGs), it at least is not a prerequisite for this action (scFv). **c** One exception to this shedding-stimulating effect was found for IgGs directed against several (repetitive) epitopes within the disordered and flexible N-terminal half of PrP (e.g. POM2; blue). Such ligands rather cause a strong surface clustering (possibly by multimeric crosslinking [as indicated in this scheme] or other structural alterations facilitating tight molecular interaction) followed by fast endocytosis and (lysosomal) degradation of PrP (which may likewise be beneficial against neurodegenerative processes).

With regard to AD, released forms or proteolytic fragments of PrP (such as N1), but likewise recPrP or PrP^C^ on extracellular vesicles, confer neuroprotection in models of AD [11, 41–46]. How exactly this is achieved on the molecular level is less clear to date. In cell-free systems, recPrP limits the growth of Aβ fibrils, while vesicle-bound PrP^C^ accelerates the formation of these supposedly less toxic Aβ species [67, 110]. In analogy to our finding for prion diseases, data obtained by using a murine AD model and our sPrP^G228^ antibody speaks in favor of a (likely beneficial) role of sPrP in promoting Aβ fibrillization and plaque formation, possibly at the expense of mobile neurotoxic AβO in the extracellular space. In fact, we show that – in analogy to what we observed in prion-infected mouse brain – sPrP is completely redistributed to particular deposits in brains of 5xFAD mice, where it is likely bound to Aβ and seen in the center of many amyloid plaques. Similar effects were also observed in AD patients, where PrP localizes to dense cores within Aβ plaques and it was suggested (though not proven) that these may represent “released” forms of PrP^C^ [68]. This reconciles data showing that PrP^C^ in its plasma membrane-bound state acts as a mediator of AβO neurotoxicity [19, 111–114], yet when overexpressed as a GPI-anchorless version rather blocks neurotoxicity [41]. Our data also suggest that it is in fact sPrP, which is responsible for the plaque-promoting effect initially ascribed to PrP^C^ “in general” in earlier studies [68–71]. Thus, physiologically sPrP may indeed act protective in prion diseases and AD by blocking toxic oligomers and/or by precipitating these into less toxic deposits (Fig. 8A). Though not assessed here, it appears likely that sPrP may act similarly against other proteinopathies, given the central role of PrP^C^ as neuronal toxicity receptor [11, 14, 18]. It would also be interesting to assess whether sPrP bound to such oligomers and deposits may serve as an “eat-me” signal for internalization by phagocytic cells [115]. In sum, stimulation of PrP^C^ shedding may be a promising therapeutic target in neurodegenerative diseases, yet further studies on this are clearly required.

In an effort to identify inducers of the ADAM10-mediated PrP shedding, we used a candidate approach and took advantage of known principles of an increased shedding caused by antibodies directed against some other ADAM10 substrates [49, 50]. This has for instance been shown for CD44, where antibody binding to this protein leads to increased release [51]. We found that several PrP^C^-directed antibodies and derivatives significantly increase its ADAM10-mediated shedding. In order to investigate the underlying mechanism of this, we assessed if antibody-induced dimerization (and hence longer retention) of PrP^C^ at the cell surface is a prerequisite. For the ADAM10 substrate CD44, dimerization is required for increased shedding [51]. Moreover, some PrP-directed antibodies indeed cause PrP^C^ crosslinking [116], and forced dimerization of PrP^C^ leads to increased proteolytic processing, mostly by elevated α-cleavage of PrP^C^ [78]. Although we cannot rule out that dimerization contributes to increased shedding upon binding of fl-IgGs to PrP^C^, the fact that we also found increased sPrP levels upon treatment with PrP-directed single-chain antibodies rather demonstrates that crosslinking is at least not a general prerequisite for the sheddingstimulating effect of such ligands. A related potential explanation for increased shedding could be antibody-mediated changes in steady-state levels of PrP^C^ at the plasma membrane, the place where ADAM10 and PrP^C^ meet and where shedding occurs. Indeed, PrP^C^-binding IgGs (exemplified here for POM1) lead to increased cell surface levels of PrP^C^, yet again, single-chain antibodies, which basically seem as effective as their full-length counterparts in their ability to stimulate shedding, do not do this. Thus, as above we conclude that altered PrP^C^ plasma membrane levels may contribute to increased shedding, yet are not a prerequisite for this. Regulation of ADAM10 substrate specificity and sheddase activity is complex, poorly understood, and may involve conformational changes of the substrate [117]. In order to investigate if a ligand-induced conformational change within PrP^C^ occurs upon binding of an antibody, we took advantage of small-angle X-ray scattering (SAXS) and focused on antibody-induced structural transitions of PrP^C^ that could modify its interaction with and support subsequent cleavage by ADAM10. Our data using recPrP suggest that the highly flexible N-terminal tail of PrP is indeed relatively moved away from the C-terminal half (and likely the GPI anchor of PrP^C^) upon antibody binding. This might support increased exposure of the very C-terminal part (containing the cleavage site), previously ‘shielded’ by movements [80] or intramolecular and possibly even described plasma-membrane interactions of the N-terminal tail [81–84], and favor access of the membrane-proximate catalytic domain of active ADAM10. This is in agreement with publications showing that Fab fragments binding to similar PrP domains influence its conformation by preventing intramolecular interactions [80] and data proposing such interactions of N- with C-terminal domains of PrP^C^ in the absence of antibody binding [84].

Lowering PrP^C^ amounts clearly is among the most promising strategies for causal treatment of prion diseases [27, 95, 96]. In fact, there is a linear inverse correlation between PrP^C^ expression levels and susceptibility towards prion diseases [22] with a complete lack of susceptibility when PrP^C^ is absent [21]. Reduction of PrP^C^ levels even has therapeutic potential once the disease becomes clinically apparent [24]. However, as with many complex diseases, combination therapy targeting different pathomechanistic aspects will likely be superior over a therapeutic strategy focussing on one mechanism only. Even if current strategies to lower total PrP expression succeed, additional stimulation of PrP shedding may add further protection (by generating sPrP as a potential anti-prion agent) while preserving certain physiological functions potentially carried out by released PrP fragments (such as myelin maintenance in the periphery [39, 106]). Interestingly, the POM2 antibody was shown to be neuroprotective and this was linked to impairing harmful interactions of the flexible N-terminal tail of PrP with the plasma membrane occurring in prion disease [81, 83, 84]. Here, we show that POM2, due to its unique binding characteristics (partially shared with POM11) and in contrast to other ligands tested here, leads to strong multimeric clustering of PrP^C^ at the plasma membrane and subsequent cellular uptake of these clusters for lysosomal degradation (Fig. 8C). This reconciles above-mentioned concepts of lowering PrP levels and implies that, in addition to POM2’s role in preventing toxic PrP N-terminus-to-membrane interaction, it efficiently causes removal of PrP^C^ from the cell surface and a reduction in total PrP^C^ levels. Since administration of antibody would allow to do this in a reversible and potentially controllable fashion, therapeutic applications are also conceivable here.

As introduced in detail above, anti-PrP antibodies have been proposed as potential prion and AD therapeutics (reviewed in [73, 118]) and, in fact, in a number of studies anti-PrP antibodies led to delay of disease onset and reduced signs of neurodegeneration [59, 65, 119, 120]. Although details are not completely understood, for prion diseases this beneficial effect is generally attributed to reduced PrP^Sc^ replication thought to be caused by either direct sterical hindrance of the critical PrP^Sc^-to-PrP^C^ interaction, stabilization of the native conformation and/or altered cellular trafficking of PrP^C^ [53, 54, 121]. For AD, binding of antibodies to membrane-bound PrP^C^ is likewise thought to block the interaction with toxic AβO species and, thus, to restrict respective surface PrP^C^-dependent synaptotoxic signaling cascades [122]. Our finding that the same anti-PrP antibodies, which have been used in some of the studies mentioned above, also induce the ADAM10-mediated shedding of PrP^C^ (Fig. 8B) adds another level of complexity and potential explanations to these data and may indicate that stimulated shedding raising extracellular amounts of seemingly neutralizing sPrP –at least in part– accounts for the protective effects.

For prion diseases, it is conceivable that sPrP, similar to what was shown for a secreted PrP dimer [40], contributes to stopping the buildup of PrP^Sc^ by binding to critical seeds thereof, thus acting dominant-negative against the misfolding of cell-associated PrP^C^. This is only seemingly in conflict with data from transgenic mice showing that anchorless PrP readily misfolds into PrP^Sc^, which is then preferentially deposited as plaques [123–125]. So, how does physiologically produced sPrP differ from the highly misfolding-prone anchorless PrP? Although this certainly deserves further investigation, a likely explanation may lie in the apparently different glycosylation state. While physiological sPrP is preferentially fully (i.e., di-) glycosylated [29], transgenically expressed secreted PrP is mainly un- and monoglycosylated. Fittingly, for many prion strains (such as the RML isolate used here), diglycosylated PrP is a relatively poor substrate [126–129], which is also reflected by relative differences in the glycopattern between PrP^C^ and PK-digested PrP^Sc^ in brain samples. Moreover, earlier cell culture experiments, where a broad range of anti-PrP antibodies inhibited PrP^Sc^ replication, revealed that ‘PrP levels’ in the media were increased [121]. According to the data presented here, this most likely represents sPrP. In the case of AD, it may well be that anti-PrP antibodies contribute to mitigating neuroprotection threefold, namely (i) by directly blocking interaction with toxic AβO species [122], (ii) by removing PrP, the receptor for toxic AβO [111] from the plasma membrane, and (iii) by hence increasing levels of sPrP, which in turn precipitates toxic AβO into possibly less-toxic fibrillary species as described earlier for recPrP or the PrP-N1 fragment. In the latter regard and to eventually develop therapeutic approaches, it will be important to further explore ligands increasing the shedding without occupying critical PrP binding sites for toxic conformers.

We found that single-chain anti-PrP antibodies efficiently induce the ADAM10-mediated shedding of PrP. Thus, not only large antibodies, but also smaller molecules resembling PrP-ligands are able to induce shedding (Fig. 8B). On the other hand, the four small chemical compounds described to bind PrP and tested here, did not induce increased release. The latter may fit a recent report suggesting that PrP is a difficult-to-target protein, at least with small ligands, and implying that, for PrP ligands to be therapeutically effective, they have to have a certain size and binding characteristics [130]. This, in turn, reinforces the use of nonsmall-molecules, such as antibodies or their derived smaller fragments, which is currently pursued despite obvious difficulties in administration and pharmacokinetics [74]. Our finding of shedding-stimulating ligands is also of interest in two different perspectives. Firstly, it may give hints to further physiological functions of PrP, which upon ligand-binding and release may act as a ligand itself serving intercellular communication, possibly in complex with the bound partner protein. It will thus be up to future studies to look for endogenous PrP ligands or interacting partners promoting its shedding. Using our sPrP-specific antibody in appropriate paradigms, this should be a feasible task. Secondly, it opens up the possibility to screen for or custom-design novel anti-PrP ligands promoting the shedding, which may then be used in therapeutic settings. Given the fact that these ligands could potentially be smaller and pharmacokinetically more favorable than antibodies, this could be an attractive aim for future studies on new therapeutic options against prion diseases and other neurodegenerative proteinopathies alike.

## Acknowledgments

We thank the Mouse Pathology Core Facility (Kristin Hartmann), the FACS Sorting Core Unit, and the Microscopy Imaging Facility (UMIF) (all at UKE Hamburg) for excellent technical support, Dr. Camilla Giudici and Dr. Michael Willem (LMU Munich) for providing N2a PrP-/- cells, and Giovanni Spagnolli (University of Trento) for the design of Suppl. Figure 6A. Molecular graphics and analyses were performed with UCSF Chimera, developed by the Resource for Biocomputing, Visualization, and Informatics at the University of California, San Francisco, with support from NIH P41-GM103311.

## Data and materials availability

All data needed to evaluate the conclusions in the paper are present in the paper and/or the Supplementary Materials. The SAXS models and the corresponding data have been submitted to the SASBDB (www.sasbdb.org), with accession codes SASDLT2 (6D11 IgG monoclonal antibody) and SASDLU2 (1:2 ratio complex between 6D11 IgG and recombinant murine prion protein). Additional data and materials related to this paper may be requested from the authors.

## Material & Methods

### Animals and ethics statement

Mice and rats were housed at standard laboratory conditions (12 hours’ light/dark cycle, constant room temperature and humidity, with food and water access ad libitum) in authorized animal facilities at Christian Albrechts University Kiel, Georg August University Göttingen, University of Zurich, and University Medical Center Hamburg-Eppendorf (UKE). Breeding, sacrification for the sake of organ removal (slice cultures, primary neurons) and all experimental procedures were approved by the respective ethical research committees of local German, French or Swiss authorities *(Freie und Hansestadt Hamburg – Behörde für Gesundheit und Verbraucherschutz* (permit numbers 48/09, 84/13, ORG587, ORG739); *Schleswig-Holsteinisches Ministerium für Energiewende, Landwirtschaft, Umwelt und ländliche Räume,* Kiel (V241-25481/2018(30-3/16); Federal state of *Niedersachsen* (permit 13/1232); *Comité d’éthique pour l’expérimentation animale,* Paris (permits Ce5/2012/018 and #1339 – 2015073113467359 v4); and the Veterinary office of the Canton of Zurich (permit ZH236/19)). Procedures were in agreement with the principles of good laboratory animal care (NIH publication No. 86-23, revised 1985) and the respective rules in the Guide for the Care and Use of Laboratory Animals of the German Animal Welfare Act on protection of animals. 5xFAD/*Prnp*^0/0^ mice were obtained by crossing *Prnp*^0/0^ mice with 5xFAD mice and crossed back with *Prnp*^0/0^ for at least 6 generations (mixed 129/Sv and C57BL/6J background). As control mice we used 5xFAD mice crossed back with WT mice for at least six generations (mixed 129/Sv and C57BL/6J background). Genotyping was performed by PCR from the tail biopsy. Intracerebral inoculations of mice with the mouse-adapted prion strain RML, subsequent observation and termination were described in detail in [38].

List of mouse lines assessed in this study:

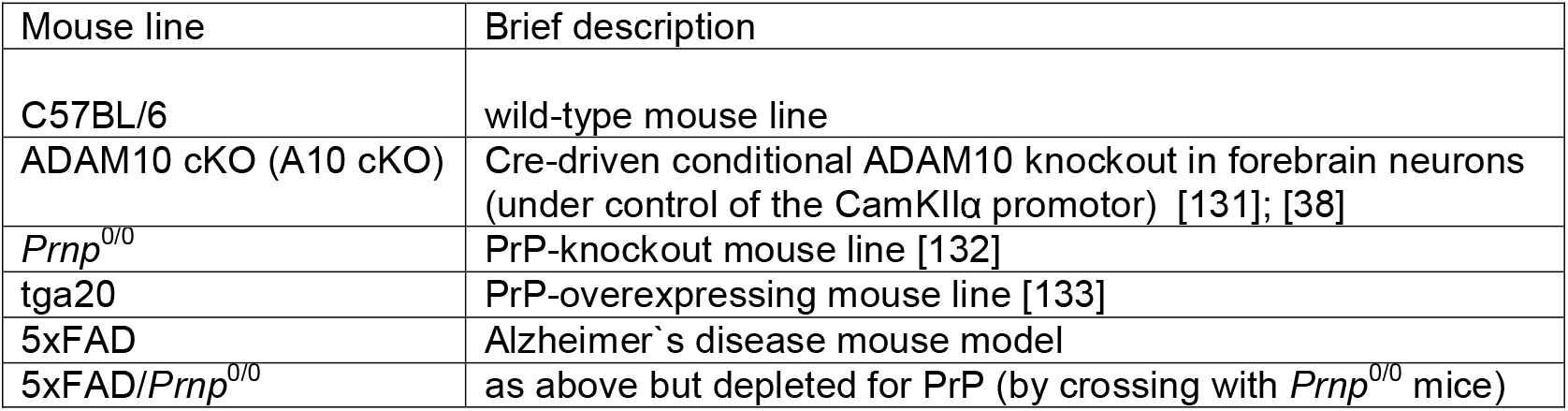

### Antibodies

List of primary antibodies used herein (incl. application):

[WB=western blot; IHC=immunohistochemistry; IF=immunofluorescence; EM=immuno electron microscopy; T=treatment; SPT-QD: single particle tracking with quantum dots]

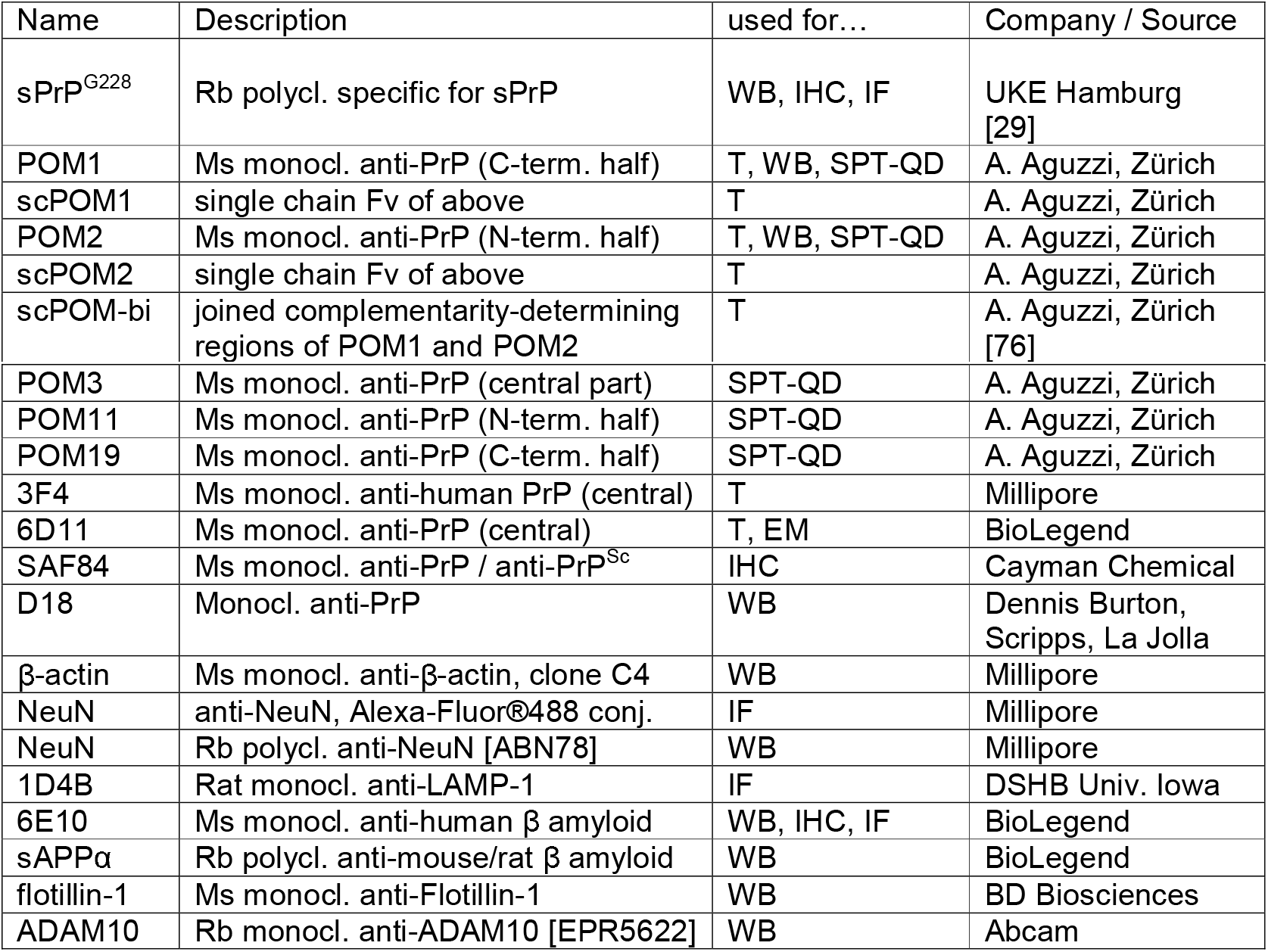

### Murine cerebellar organotypic slice cultures (COCS)

350 μm thick COCS were prepared from 9-12 day-old *Prnp*^0/0^, C57BL/6 and tga20 pups according to a previously published protocol [134]. 6-9 COCS were plated per PTFE-coated cell culture insert (Millipore) in organotypic slice culture medium [134]. Pharmacological treatment was initiated 10-14 days after dissection of COCS and was re-added with every medium change (treatment duration: 8 days with 3 media changes for Fig. 3A-C; Fig. 3D: IgG and 6D11: 14 days, STS: 3 days). Final concentration of antibodies (6D11, 3F4, pooled murine IgG) was 67nM diluted in organotypic slice culture medium. Protein analysis and immunohistochemical analysis of COCS were described extensively before [76].

### Quantitative RT-PCR

RNA was isolated from N2a cells upon antibody treatment for 18 h using the RNeasy Mini Kit (Qiagen) following the manufacturer’s instructions. 2 μg of DNAse-treated RNA was used for cDNA synthesis using the RevertAid cDNA Synthesis Kit (Thermo Fisher Scientific). The gene expression analysis for *Prnp* and *Gapdh* was performed using the respective TaqMan probes (Thermo Scientific Fisher). PrP^C^ expression levels were depicted as percentage of GAPDH expression using the ΔCT method for calculation.

### Preparation of mouse brain homogenates

Pieces of fresh or frozen frontal brain were processed as 10% (w/v) homogenates in RIPA buffer (50 mM Tris-HCl pH 8, 150 mM NaCl, 1% NP40, 0.5% Na-Deoxycholat, 0.1% SDS) with freshly added cOmplete™ protease inhibitor cocktail (PI; Roche). Tissue was ground by 30 strokes on ice using a Dounce homogenizer and subsequently incubated on ice for 15 min before resuspending by pipetting up and down. Homogenates were then spun down at 11,000×*g* for 10 min at 4°C. Resulting supernatants were collected, aliquoted and stored at −80°C. For further use 30 μL were mixed with 120 μL of ddH_2_O and 50 μL of 4× loading buffer (250 mM Tris–HCl, 8% SDS, 40% glycerol, 20% β-mercaptoethanol, 0.008% Bromophenol Blue, pH 6.8) and boiled for denaturation for 8 min at 96°C.

Prion-infected mice and samples were handled in respective biosafety facilities/laboratories at the UKE Hamburg. For detection of PrP^Sc^ in prion-infected mouse brains, 20% (w/v) homogenates of frontal brain were prepared using phosphate buffer saline (PBS) lacking protease inhibitors. Homogenates were prepared as above using a Dounce homogenizer and then centrifuged at 1500×*g* for just 2 min to pellet crude cellular debris. 4 μL of the supernatant were incubated with 20 μg/mL proteinase K (PK; Roche) in a total volume of 22 μL RIPA buffer for 1 hr at 37°C to digest all proteins except for PK-resistant PrP^Sc^. Digestion was terminated by addition of 2.5 μL of 10× loading buffer and boiling for 8 min at 96°C.

### Cell culture, treatments, lysis, harvesting of media and sample preparation

#### Preparations and treatments

Murine neuroblastoma cells (N2a; ACC148, DSMZ Germany) and the embryonic mouse hippocampal cell line mHippoE-14 (CLU198; CELLutions Biosystems Inc.) were cultured at 37°C in an atmosphere of 5% CO_2_ in Dulbecco’s modified Eagle’s medium (DMEM; Thermo Fisher Scientific) supplemented with 10% fetal bovine serum (FBS; Thermo Fisher Scientific). Generation of PrP-depleted N2a cells (*Prnp* KO) was described elsewhere [135]. For overexpression of murine wild-type (PrP-WT) or 3F4-tagged PrP (PrP-3F4), these PrP knockout cells were transiently transfected with the respective constructs [29] using Lipofectamine 2000 (Thermo Fisher Scientific) following the manufacturer’s instructions.

One day before treatments, 250,000 cells were seeded per well in 6-well plates. The next day, media was freshly exchanged to pre-warmed OptiMEM (Gibco) and treatments were started by adding antibodies and/or compounds in the stated concentration to the medium. Treatments were carried out overnight for 18 h in a total of 1 mL of OptiMEM.

#### Lysis of N2a and mHippo cells and processing of conditioned media

Following treatments, conditioned media and corresponding adherent cells were quickly yet carefully harvested in parallel working on ice or at 4°C. Media was aspirated, transferred to a 1.5 mL Eppendorf tube already containing 50 μL of a pre-dissolved and 20x concentrated PI (in PBS) and gently inverted for mixing. Cells were washed twice with cold PBS and incubated for 10 min with 150 μL RIPA buffer (with freshly added PI), scraped off from the plate and transferred to a 1,5 μL tube, shortly vortexed and incubated on ice for additional 15 min. After a centrifugation step at 12,000×*g* and 4°C for 12 min, the supernatant (=cleared lysate) was transferred to a fresh tube and either stored at −80°C or mixed (50 μL lysate + 50 μL 4x sample buffer + 100 μL ddH_2_O), boiled (10 min at 96°C) and loaded for SDS-PAGE. Harvested media supernatants were centrifuged at 500×*g* and then 5,000×*g* (both for 5 min at 4°C) and at each step 50 μL were left at the bottom (as a pellet of cellular debris was mostly invisible). The remaining 900 μL were subjected to protein precipitation as follows: 1/100 volume (i.e. 9 μL) of a 2% sodium deoxycholate (NaDOC) solution was added to the sample and mixed by vortexing followed by 30 min of incubation on ice. Samples were then mixed with a 1/10 volume (i.e. 90 μL) of 100% (6.1N) trichloroacetic acid (TCA; Sigma Aldrich) and incubated for another 30 min on ice. After centrifugation at 15,000×*g* for 15 min at 4°C, the supernatant was aspirated and the pellet air-dried for 5 min. The pellet was completely resuspended (by intensively pipetting up and down and mixing at 800 rpm at 50°C for 15 min) in 100 μL of 1× loading buffer. As the blue color changed to yellow (indicating low pH), 1.5 μL of 2M NaOH was mixed into the sample and blue color reappeared. Samples were boiled for 10 min at 96°C and stored at −80°C or directly used for SDS-PAGE.

#### Treatment of HEK293 cells with PrP-directed chemical compounds

HEK293 cells were stably transfected with a pcDNA3.1 plasmid carrying a hygromycin resistance cassette and encoding for mouse wild type PrP under the control of the CMV promoter. On day 1, cells were seeded at ~70% confluency in 24-well plates (medium containing: DMEM, FBS 10%, NEEA, PEN/Strep, L-Glutamine, 150ug/ml hygromycin); on day 2, medium was replaced with fresh medium containing each chemical compound at 0.1, 0.3, 1, or 3μM (but lacking hygromycin); on day 3, conditioned medium was collected and diluted 2:1 in 4x Laemmli sample buffer (2% SDS, 10% glycerol, 100 mM Tris-HCl pH 6.8, 0.002% bromophenol blue, 100 mM DTT), while the corresponding adherent cells were directly lysed with 2x Laemmli sample buffer.

### Cell surface protein biotinylation assay

After 18 h of treatment with antibodies, N2a cells were washed twice with cold PBS and incubated for 30 min with 0.5 mg/mL EZ-Link Sulfo-NHS-SS-Biotin (Thermo Fisher Scientific) in PBS shaking at 4°C. Cells were washed 3x for 5 min at 4°C with 0.1% BSA and lysed with 500 μL RIPA buffer. After complete lysis, the supernatant was diluted 1:1 with Triton dilution buffer (100mM TEA, 100mM NaCl, 5mM EDTA, 0.02% NaN3, 2.5% Triton X-100, pH 8.6, +PI) and incubated for 1 h with 200 μL NeutrAvidin agarose beads (Thermo Fisher Scientific) at 4°C. Beads were washed with wash buffer 3x (20mM TEA, 150mM NaCl, 5mM EDTA, 1% Triton X-100, 0.2% SDS, 0.02% NaN_3_, pH 8.6, +PI) and centrifuged at 1,000×*g*. Two additional washing steps were performed with the final wash buffer (20mM TEA, 150mM NaCl, 5mM EDTA, pH 8.6, +PI). Finally, 50 μL of 4x sample buffer (including DTT) were added and the samples were boiled for 6 min at 96°C to release and denature biotinylated proteins from the beads. The supernatants were loaded for SDS-PAGE for further biochemical analysis.

### SDS-PAGE and western blot analyses

#### SDS-PAGE and western blotting

Denatured samples (tissue homogenates, cell lysates or precipitated media) were loaded on precast Nu-PAGE 4–12% Bis-Tris protein gels (Thermo Fisher Scientific), self-cast 10% or 12% SDS-gels or Any kD™ Mini-PROTEAN® TGX™ Precast Protein Gels (BioRad). After electrophoretic separation, wet blotting (at 200 mA per gel for 1h) was applied to transfer proteins onto nitrocellulose membranes (BioRad). Membranes were then blocked for 45 min with either 1× RotiBlock (Carl Roth) in TBS-T or 5% skimmed dry milk (in TBS-T) under gentle agitation at RT. Membranes were incubated overnight with the respective primary antibodies (see list above) diluted 1:1,000 (1:2,500 for POM antibodies; 1:5,000 for β-actin) in the respective blocking reagents at 4°C on a shaking platform. The next day, membranes were washed 4x for 5 min with TBS-T and incubated for 45 min at RT with HRP-conjugated secondary antibodies. After extensive washes (at least 6x for 5 min) in TBS-T signal detection was performed (after incubating blots for 5 min with either Pierce ECL Pico or Super Signal West Femto substrate (Thermo Fisher Scientific)) with a ChemiDoc imaging station (BioRad). Densitometric quantification was done using the QuantityOne software (Biorad) followed by further analysis in Excel (Microsoft).

#### SDS-PAGE and western blotting [of compound-treated HEK293 cells]

A 30 μL aliquot of either medium or lysate for each sample was heated at 95 °C for 10 min and loaded on two separate SDS-PAGE gels (Any kD™ Mini-PROTEAN® TGX Stain-Free Protein Gels; BioRad). Proteins were electrophoretically transferred to polyvinylidene fluoride (PVDF) membranes, which were then blocked for 20 min in 5% (w/v) non-fat dry milk in Trisbuffered saline containing 0.05% Tween-20. After incubation with appropriate primary (D18 [1:5,000] and sPrP^G228^ [1:3,000]) and secondary antibodies, signals were revealed using enhanced chemiluminescence (Luminata, BioRad), and visualized by a Bio-Rad XRS Chemidoc image scanner (BioRad). Values were obtained by densitometric quantification of PrP bands using the ImageLab 5.2.1 software (ChemiDoc, Bio-Rad). Each PrP signal (D18 or sPrP^G228^) was normalized to the signal of total proteins in cell lysates (directly acquired by detecting the fluorescence of proteins in stain-free gels) and expressed as percentage of the DMSO-treated control.

### Annexin V cell toxicity assay

N2a cells were grown on 6-well plates until 80% confluency, treated overnight with antibodies, washed with PBS and detached by incubating for 5 min with accutase (Thermo Fisher Scientific) at 37°C. Next, cells were washed with PBS and resuspended in 1x binding buffer (provided by manufacturer) at 1-5 x 10^6^ cells/mL. 5 μL of PE-conjugated Annexin V (Thermo Fisher Scientific) was added to 100 μL of the cell suspension and incubated at room temperature in the dark. After 15 min of incubation, cells were again washed with 1x binding buffer and subsequently analysed by flow cytometry. PE-positive, apoptotic cells were counted by using flow cytometry in a FACSCanto II (BD Bioscience, Franklin Lakes, NJ, USA). As a positive control, cells were treated with different concentrations of staurosporine (STS), a known inducer of apoptotic cell death.

### Immunohistochemistry (IHC)

Mouse brains were dissected and immediately fixed overnight in an excess volume of 4% buffered formalin. Prion-infected samples were inactivated for 1 h in 98–100% formic acid prior to transfer into clean tubes and export from the biosafety facility. The latter samples were then again incubated for another night in 4% buffered formalin at 4°C. Formalin-fixed samples then underwent dehydration and were embedded in low melting point paraffin blocks following standard histological protocols. Sections of 3-4 μm were prepared with a microtom submitted to immunostaining following standard immunohistochemistry procedures using the Ventana Benchmark XT machine (Roche Diagnostics). In brief, sections were deparaffinated and then boiled for 30–60 min in 10 mM citrate buffer (pH 6.0) for antigen retrieval. All applied solutions were purchased from Ventana. Afterwards, sections were incubated with primary antibodies diluted in 5% goat serum (Dianova, Hamburg, Germany), 45% Tris buffered saline (TBS) pH 7.6, 0.1% Triton X-100 in antibody diluent solution (Zytomed, Berlin, Germany) for 1 hour. Primary antibodies used for IHC analyses were (for further information also refer to list above): sPrP^G228^ (1:30, for detection of sPrP), SAF84 (1:100, for PrP^C^ and PrP^Sc^), 6E10 (1:100, for human APP and Aβ deposits). Secondary antibody treatment was performed using anti-rabbit histofine Simple Stain MAX PO Universal immunoperoxidase polymer or Mouse Stain Kit (for detection of mouse antibodies on mouse sections); all purchased from Nichirei Biosciences (Tokyo, Japan). Final visualization of antibodies was achieved with the Ultra View Universal DAB Detection Kit (brownish signals) or Ultra View Universal Alkaline Phosphatase Red Detection Kit (yielding pink signals) from Ventana using standard machine settings. Notably, whenever possible experimental groups were stained in one run, thus ensuring identical conditions and good comparablity. Light blue counterstaining was likewise done by the machine following standard procedures. Secondary antibody-only controls were always run when establishing a given staining yet never revealed a relevant signal. For PrPSc detection using SAF84 antibody, sections of 4 μm were pretreated with 98% formic acid for 5 min. Moreover, sections were boiled in 1.1 mM sodium citrate buffer (2.1 mM Tris-HCl, 1.3 mM EDTA, pH 7.8) at 95°C for 30 min, then incubated with PK for 16 min, in Superblock for 10 min, finally incubated with primary antibody and further processed as above.

Stained sections were assessed and representative pictures were taken in TIF format on a digital microimaging device (DMD108, Leica). Pictures were further processed by performing a white balance against a negative, non-stained area within the same field, and finally assembled (using fixed/scaled aspect ratio) for comparison of experimental groups using Photoshop (Adobe).

### Immunofluorescence (IF) stainings and (live) microscopy

#### IF stainings of free-floating murine brain sections and microscopic assessment

Mice were deeply anaesthetized mice with Rompun/Ketamin followed by transcardial perfusion with 0.1M phosphate buffer (PB; pH 7.4) and consecutively 4% paraformaldehyde (PFA). Brains were removed and post-fixed by immersion. After 4 hours of post-fixation, PFA was removed and replaced by 30% sucrose (w/v) in 0.1M PB. After incubation overnight, brains were sectioned sagittally at 35 μm thickness as free-floating sections with a Leica 9000s sliding microtome (Leica, Wetzlar, Germany). For immunofluorescence staining, the sections were blocked in blocking solutions (0.5% Triton-X 100, 4% normal goat serum in 0.1M PB, pH 7.4), incubated in blocking solution containing the primary antibodies at 4°C overnight. After three washes with wash solution (0.1M PB, pH 7.4 containing 0.25% Triton-X 100), sections were incubated for 90 minutes with secondary antibody in solution, washed again three times in wash solution containing 4’,6-Diamidin-2-phenylindol (DAPI), finally brought on glass slides and embedded in Mowiol/DABCO (1,4-diazabicyclo-[2.2.2]-octane). Images were acquired with an Olympus FV1000D Laser Scanning Confocal Microscope (model: FV10-292-115) with a 60x lens (UPLSAPO) and processed with the FV10-ASW 4.2 ViewerSoftware (Olympus, Germany).

#### IF analyses and (live) microscopy of antibody-treated rat primary neurons

Primary hippocampal cultures were prepared as described previously [136] from E18 Wistar rats Unilever HsdCpb:WU (Envigo) and plated at a density of 60,000 cells per coverslip in a 12-well plate.

For live imaging experiments, neurons were transfected with eGFP-tagged PrP [cloning information see below] (1μg DNA per well) at day in vitro (div) 14 using Lipofectamine 2000 according to the manufacturer’s instructions. At div15 live imaging of transfected neurons was performed at a Nikon Eclipse Ti microscope Ti-E controlled by VisiView software. Illumination was achieved by a 488 nm excitation laser, coupled to a CSU-X1 spinning disk unit via a single-mode fiber. Emission was collected through a quad band filter (Chroma, ZET 405/488/561/647m) on an Orca flash 4.0LT CMOS camera (Hamamatsu). Imaging in confocal mode was done with a step size of 0.5 μm and pixel size of 108.3 nm (using the 60x 1.4NA objective (Nikon, CFI Plan Apo Lambda Oil)). Neurons of similar PrP-GFP expression rate (judged by signal intensity) were imaged in 500 μL medium at basal conditions for 5 minutes and upon administration of 1 μg antibody (POM2, 6D11, 3F4) for 10 minutes with 1 image per minute, up to 1 h with fewer imaging time points. For image representation, maximum projections were created and brightness/contrast adjusted per cell.

For staining of endogenous prion protein, hippocampal cultures at div19 were treated with 1 μg POM2 antibody in 500 μL culturing medium for 30 min, and immediately afterwards fixed in 4% PFA / 4% sucrose for 10 min at room temperature. Immunocytochemistry was performed as in [136] using POM2 antibody (1:250) followed by a secondary anti-mouse-Alexa Fluor 488 antibody. Actin staining was obtained by using phalloidin-647N (1:100 dilution) in PBS and incubation for 4 h at room temperature. Images were acquired at a Leica TCS SP5 microscope (Leica microsystems), controlled by Leica Application Suite Advanced Fluorescence software. Excitation of fluorophores was done with an Argon 488 nm laser and a HeNe 633 nm laser, and signals detected on HyD detectors. Neurons were imaged using a 63x objective (1.4 HCX PL APO CS), at 400 Hz and frame averages of 2 and z-steps of 0.5 μm (overview) or 0.3 μm (dendrite). Picture format was set to 8 bit, and for the overview image to 1024 px, using no zoom (pixel size: 240 nm) and for dendrites to 5x zoom at 512 px (pixel size: 96 nm). For image representation, average projections were created and brightness/contrast adjusted per cell.

##### eGFP-tagged PrP

The moPrP-eGFP_39/40 fusion construct carrying eGFP within the flexible N-terminal tail between aa 39 and 40 of mouse PrP was created using NEB® Golden Gate Assembly (New England Biolabs). In brief, three fragments (PrP1-39, eGFP lacking initial MET and terminal STOP, PrP40-254) were amplified via PCR using oligonucleotides as primers in respective combinations (sequence provided as 5’–3’): *PrP1-39:* SP_PrP-fwd (TTTTAAGCTTATCAGTCATCATGGCGAACCTTGGCTAC) + BsaI-PrP39-rev (TGGTCTCTTCACGGGATACCGGCTTC); *eGFP*: BsaI-GFP-fwd (AGGTCTCTG TGAGCAAGGGCGAGGAG) + BsaI-GFP-rev (TGGTCTCTCTTGTACAGCTCGTCCATGC); *PrP40-254:* BsaI-PrP40-fwd (AGGTCTCTCAAGGGGCAGGGAAGCC) + pcDNA3.1-F1-rev (AGGGAAGAAAGCGAAAGG). PCR products were fused to a full length construct using a standard Golden Gate Assembly protocol (1x T4 Ligase Buffer, 500 U T4 DNA Ligase, 15 U BsaI-HF®v2, respective fragments, 30 cycles of two step incubation in thermocycler: 37°C for 5 min followed by additional 5 min at 16°C). The resulting construct was cloned into an existing pcDNA3.1-moPrP plasmid using the moPrP endogenous restriction sites AgeI and BstEII and T4 DNA ligase. Cloning success was verified by sequencing.

### Single Particle Tracking using Quantum Dots (SPT-QD)

SPT-QD studies were performed on hippocampal neurons prepared from Sprague-Dawley rat embryos (Janvier Labs, France). The SPT-QD protocol and analysis methods have been used and described in several previous publications [137–139]. POM-x (POM1, POM2, POM3, POM11 or POM19) exposure was performed on live neurons in the culture medium maintained in an incubator at 37°C and 5% CO_2_. MEM recording medium (Phenol red-free MEM, 33mM glucose, 20mM HEPES, 2mM glutamine, 1mM sodium pyruvate, and 1x B27) was used for Quantum dot labeling and imaging. Neurons were labeled using POM-x antibody pre-coupled with QD-605 nm. To pre-couple antibodies with QDs, the following protocol was applied: Mix 1 μL POM-x antibody + 1 μL Fab’-QD-605 + 7 μL 1x PBS and gently shake for 30 min followed by addition of 1 μL 1x casein solution for additional 15 min [137]. Synapses were labeled and identified using FM4-64 labeling. SPT-QD experiments were performed on neurons aged 20-21 div. Tracking and analysis was performed using SPTrack_v4, homemade software in Matlab (MathWorks) [137]. First, the center of the QD fluorescence spot was determined by Gaussian fit with a spatial resolution of 10-20 nm. Trajectories were generated by associating spots in a given frame with spots in the previous/next frame based on a maximum likely approach. Trajectories with a minimum length of 15 consecutive frames were used and trajectories below 15 points were excluded. The mean square displacement (MSD) was calculated using MSD(ndt) = (N - n)^-1^ Σ _i_=_1_ ^N-n^ [(x_i+n_ - x_i_)^2^ + (y_i+n_ - y_i_)^2^], with x_i_ and y_i_ being the coordinates of an object on frame I, *N*: total number of steps in the trajectory, *dt*: time between two successive frames, and *ndt*: time interval over which displacement is averaged [97, 140]. MSD plot was generated and used to compute diffusion coefficient *D* by fitting the first two to five points of the MSD plot versus time with the equation MSD(t) = 4*D*_2-5t_ + 4σ_x_^2^, with σ_x_ representing the spot localization accuracy in one direction.

### Small angle X-ray scattering (SAXS) – sample preparation, data acquisition and analysis

The recombinant mouse prion protein 23-230 (recPrP) was obtained following a previously reported protocol [141]. After lyophilization, recPrP was dissolved in a buffer containing 150mM NaCl, 10mM HEPES (pH 7.5). The recPrP solution was subjected to centrifugation at 10,000 x *g* at 4°C and the supernatant was used for downstream preparation after the concentration determination. The 6D11 antibody (in PBS) was measured in a concentration series (0.75, 1, 1.5, 2 mg/mL) in the same buffer. RecPrP and 6D11 IgG were mixed in a 2:1 molar ratio to obtain the concentration series for the recPrP/6D11-antibody complex (i.e., 0.3 mg/mL, 0.5 mg/mL, 1.5 mg/mL and 1.8 mg/mL). After mixing recPrP and 6D11, the mixtures were incubated for 1 h on ice to achieve maximum recPrP-to-6D11 binding.

Synchrotron SAXS data was acquired on the EMBL beamline P12 [142] at the Petra III storage ring of DESY (Hamburg, Germany). The data were obtained using an automatic sample changer. To avoid radiation damage, the samples were exposed under a continuous flow in a quartz capillary (0.9 mm or 1.7 mm diameter). The parameters of data collection are shown in Suppl. Table 1. The scattered intensity was calibrated to absolute units using the scattering of water at 293 K.

Initial data processing and analysis was carried out using PRIMUS [143], and overall parameters were calculated using ATSAS suite [144]. A previously described SAXS data set and corresponding CORAL model of the murine recPrP data was utilized from Small Angle Scattering Biological Data Bank (SASBDB; www.sasbdb.org) accession code: SASDHV9. The modelling for the 6D11 antibody was performed with CORAL, a program allowing for the representation of models of partially disordered proteins as quasi-random loops [143, 145].

The recPrP/6D11-antibody complex was interactively modelled in SASpy [146] by optimizing the conformation of a 2:1 complex between a representative CORAL model of the antibody and an extended conformation of recPrP. The antibody Fab domains were placed to maintain contact with the epitope on the extended N-terminal part of recPrP (residues 94-109; as reported by the manufacturer). UCSF Chimera [147] was used for model depiction. Additional information is provided in association with Suppl. Table 1.

### Solubility/precipitation assay

Quantification of soluble mouse PrP and antibodies (POM1 and POM2-IgG) was performed using SDS-PAGE following a published protocol [76]. The mPrP:Ab complexes were formed in a 2:1 ratio at a concentration of 10μM and 5μM in 20mM HEPES pH 7.2, 150mM NaCl.

After complex formation, samples were centrifuged for 2 minutes at 20,000×*g* at 4°C and supernatants were collected and loaded on polyacrylamide gels (4% Stacking ± 12% Running). Proteins on gels were then acquired using Fusion FX (Vilber) according to standard procedures and quantification of soluble proteins was then performed using ImageJ, normalizing all samples to mPrP or antibodies alone. Three different time points after complex formation were analyzed: t0, t10’ (after 10 minutes) and t60’ (after 60 minutes).

### Atomic force microscopy (AFM)

Equal volumes of 10μM recPrP (in 100mM HEPES, 0.5% Triton X-100) and 5μM anti-PrP antibodies POM1 or POM2 (each in 100 mM HEPES, 0.5% Triton X-100), were separately incubated at room temperature for 1 h, with gentle shaking. Freshly cleaved muscovite mica (001) substrates were coated with abovementioned dissolution buffer only, recPrP-POM1 preparation or recPrP-POM2 preparation at a surface density of 25 μL/cm^2^. The coated mica substrates were then allowed to dry under gentle vacuum and were used for atomic force microscopy. A Nanoscope® IIIa Multimode AFM (Bruker Cooperation, Billerica, MA, USA) was used for surface characterization of PrP-POM aggregates. Data evaluation image representation was performed using WSxM 5.0 software [148]. The scans were made over the scan ranges of 5×5 μm2 and 1×1 μm^2^. Tappingmode NSL-20 probes were obtained from Nanoworld Holdings AG (Schaffhausen, Switzerland). The probes had an average force constant of 20 Nm^-1^ and were driven to oscillate at 200 ± 5 kHz for AFM imaging.

### Immunogold-labeling and electron microscopy

N2a cells were treated with POM2 or POM1 (4 μg/mL) for 5 min at 37°C. Then they were prepared, sectioned and labelled according to [149]. Briefly, cells were fixed in a mixture of 4% paraformaldehyde (PFA) and 0.1% glutaraldehyde (GA) and covered with 1% w/v gelatin in PBS after several washing steps. They were scraped and transferred into Eppendorf tubes and spun down. The pellet was resuspended in 12% w/v gelatin in PBS and solidified on ice. Small pieces of the gelatin-embedded cells were cut and blocks were left in cold 2.3M sucrose overnight. Cryoprotected pieces were mounted on specimen holders immersed in liquid nitrogen and ultrathin sections (70 nm) were cut and collected on Formvar/Carbon coated grids (Science Services GmbH, Germany). Sections were labelled either with POM2 or 6D11 antibody (dilution 1:25) and recognized with 10 nm large protein A-coupled gold (purchased from G. Posthuma, University Medical Center Utrecht). Ultrathin sections were examined in an EM902 (Zeiss, Germany). Pictures were taken with a TRS 2K digital camera (A. Tröndle, Moorenweis, Germany).

### Statistical analyses

Applied statistical tests and consideration of significance as indicated in figure captions.

## Supplementary material

**Suppl. Fig. 1:** Immunohistochemical assessment of sPrP and (human) Aβ/APP in different brain regions of 5xFAD and 5xFAD/*Prnp*^0/0^ mice. Representative pictures showing areas of dentate gyrus, subiculum and cerebral cortex. Prominent Aβ plaques as well as cell-associated Aβ/APP are detected with the 6E10 antibody (DAB) in both genotypes, whereas (plaque-like) sPrP (using the sPrP^G228^ antibody) is only detected in 5xFAD mice with WT background (i.e., with PrP expression). This excludes unspecific binding of sPrP^G228^ (and/or the respective 2^nd^ antibody used for detection) to Aβ deposits.

**Suppl. Fig. 2:** Western blot analysis of a surface biotinylation assay (**a**) showing membrane levels of ADAM10 and PrP (flotillin shown as loading control) and corresponding total lysates (**b**) showing ADAM10 and PrP amounts (actin served as loading control) in N2a cells treated with POM2 or POM1 compared to cells treated with a non-PrP-directed control antibody. Quantification in **c** shows the relative levels of PrP in lysates (PrP/actin ratio) and at the cell surface (PrP/flotillin ratio; dotted graphs) with the respective control treatment set to 1. Note the decrease in PrP upon POM2 treatment, which is particularly pronounced at the cell surface. POM1 instead caused elevated PrP levels at the plasma membrane. Statistical analysis was carried out with n=3 for all experimental groups. For comparing cell surface PrP levels to the levels of PrP in lysates in each treatment, significance values were obtained by implementing one-way ANOVA followed by the Tukey post-hoc test. Plotted data shows mean ± SEM. *p < 0.05, **p < 0.01.

**Suppl. Fig. 3:** Assessment of toxic effects of antibody treatments using an Annexin V apoptosis assay. Upon treatment with the indicated antibodies, the percentage of Annexin V-positive cells was determined by FACS compared to untreated (untr.) or 3F4 IgG-treated negative controls (grey bars). Different concentrations of staurosporine were used as positive controls inducing cell death. Mean ± SE; n=3 independent experiments (except for for 250nM STS: n=2 due to an experimental outlier excluded from quantification).

**Suppl. Fig. 4:** Antibody-mediated effects on PrP shedding in the neuronal cell line mHippo E14. To confirm findings in N2a cells we performed the same treatments in another murine brain-derived cell line. **a** Incubation with antibodies did not alter overall cell density or morphology compared to untreated controls, thus suggesting lack of major toxicity (scale bar = 50 μm). **b** Representative western blot analysis showing similar changes caused by antibody treatment as observed in N2a cells (Figure 2C). However, while increased shedding was again observed upon 6D11 or POM1 treatment, the reduction of total PrP caused by POM2, albeit detectable, was not significant after quantification of n=3 independent experiments (graphs represent mean ± SEM) shown in **c**. Graphpad Prism (6.01) was used for calculating statistical significance in multiple comparisons by one-way ANOVA and uncorrected Fisher’s LSD test (*p < 0.05).

**Suppl. Fig. 5:** SAXS profiles for the concentration series of (**a**) 6D11 antibody and (**b**) recPrP (23-230) / 6D11 antibody complex (in a 2:1 ratio). **c**, **d** SAXS data and modelling of the 6D11 antibody. **c** Experimental SAXS profile (symbols) and CORAL fit (solid line) for the representative model of the 6D11 antibody (χ^2^=0.85). **d** A typical model of the 6D11 antibody obtained with CORAL from the IgG2a crystal structure 1igt.pdb allowing for a flexible hinge region (the model was further used for the modelling of the recPrP/6D11 complex shown in figure 4). For further details also refer to Suppl. Table 1 and its associated information.

**Suppl. Fig. 6:** No shedding-stimulating activity of four PrP-directed chemical compounds. **a** Structural representation of four known PrP-binding compounds (framed boxes) and their expected binding regions (coloured regions) within the globular part of PrP (center). **b** Biochemical assessment of total proteins (as loading control) and PrP levels in lysates as well as sPrP in corresponding media supernatants of HEK293 cells stably expressing murine PrP and treated with increasing concentrations of the respective substance (indicated). Densitometric quantifications of at least four independent experiments are shown under the representative blots. PrP signals (detected with D18 or sPrP^G228^ antibody) were normalized to the signal of total proteins in cell lysates. Bar graphs show PrP or sPrP levels expressed as the mean percentage of untreated (DMSO only) controls ± standard errors. Data was processed with the Prism software, version 7.0 (GraphPad) and analyzed with one-way ANOVA test, Dunnett’s post-hoc test; *p*-values are indicated as **<0.01, ***<0.001.

**Suppl. Fig. 7:** Fluorescence microscopy demonstrating POM2-mediated clustering of PrP. **a** Live-microscopy pictures (see also Suppl. Movie 1) of rat hippocampal neurons expressing GFP-tagged PrP (green) taken shortly before (left two columns; −5 and 0 min) and after (right three columns; 5, 10 and 60 min) treatment with POM2, 6D11 or 3F4 antibody (start of treatment indicated by red arrow). Overviews showing neuronal dendritic trees (scale bars = 20 μm) are followed by magnified view on individual dendrites indicated by white frames (scale bar in close-ups is 4 μm). Note that a strong clustering of PrP (green) is only detectable upon treatment with POM2. **b** IF analysis of WT rat neurons fixed after 30 min of POM2 treatment and stained for PrP (green) reveals that POM2-mediated clustering also occurs on endogenous PrP. Phalloidin (red) was used to stain actin for better display of neuronal processes.

**Suppl. Fig. 8:** Immuno-EM comparison between N2a cells treated for 5 min with either POM1 or POM2. Immunogold-positive clusters were exclusively observed in samples incubated with POM2 (lower panel), whereas treatment with equal amounts of POM1 only resulted in isolated dots (upper panel). In this set of experiments, pan-PrP antibody 6D11 was used for detection, followed by a rabbit anti-mouse secondary antibody and protein A coupled to 10 nm colloidal gold. Ex: extracellular; in: intracellular.

**Suppl. Movie 1:** Time-resolved compilation of a representative live microscopy analysis of PrP-GFP-expressing rat neurons treated with POM2 (left), 6D11 (center) or 3F4 (right) antibodies (parts of it are highlighted in Fig. 6 and Suppl. Fig. 7A). Note the strong clustering of PrP upon POM2 administration visible all over the selected dendritic tree.

**Suppl. Table 1:** SAXS data collection, parameters and additional information.

**Suppl. Table 2:** SPT-QD analysis.

