## Supplementary figures and images for "Ligands binding to the cellular prion protein induce its protective proteolytic release with therapeutic potential in neurodegenerative proteinopathies"

### Suppl Figure 1

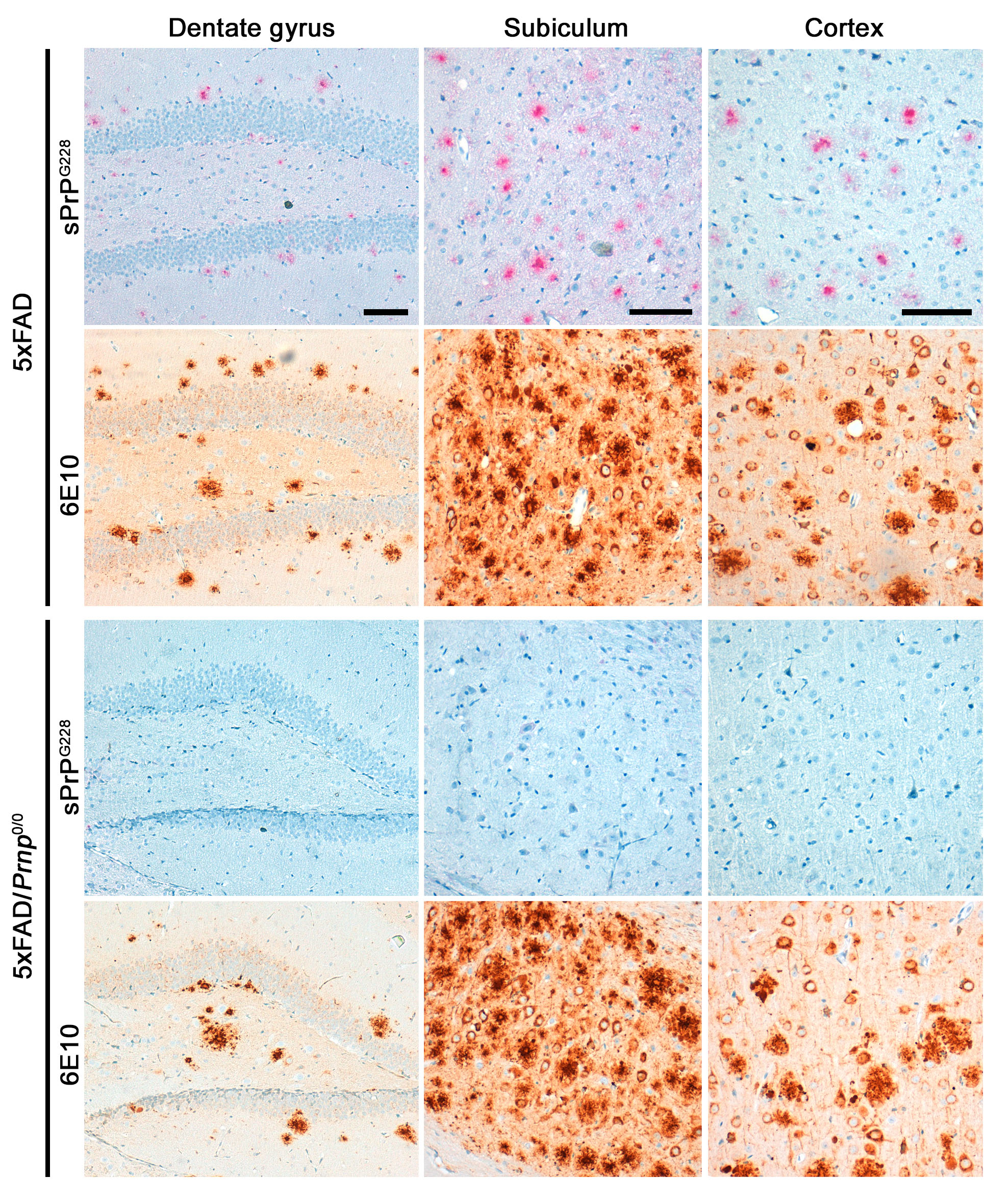

### Suppl Figure 2

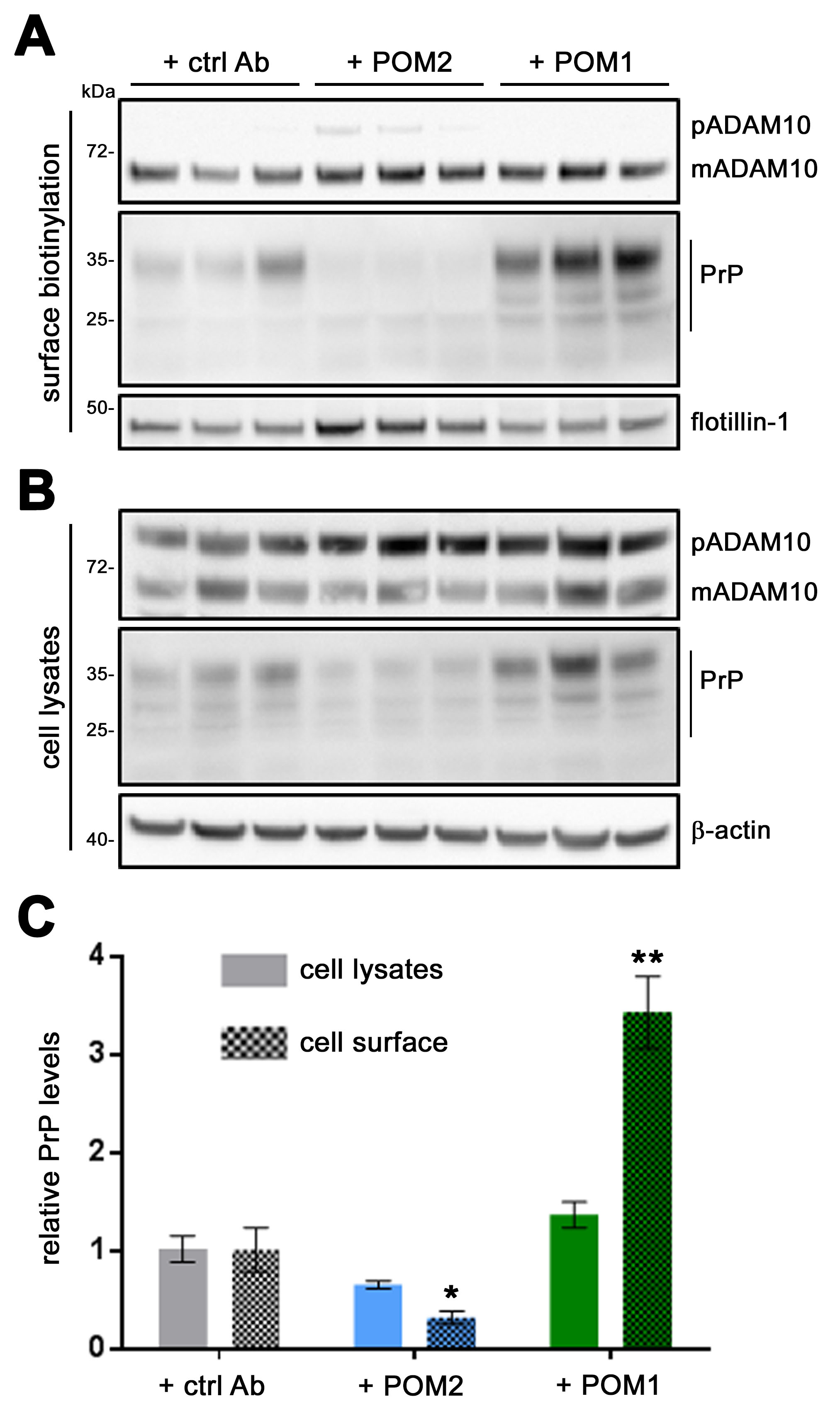

### Suppl Figure 3

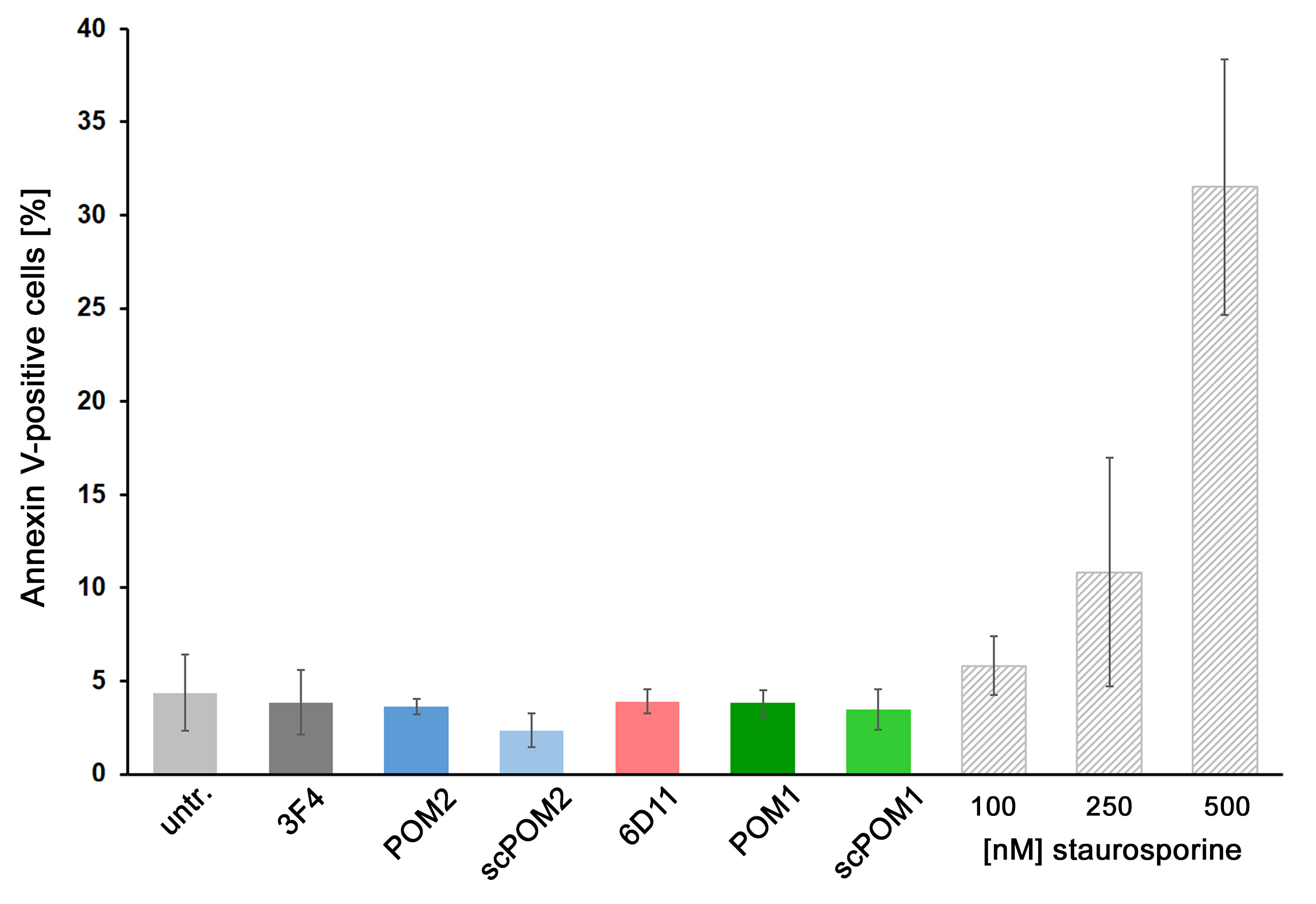

### Suppl Figure 4

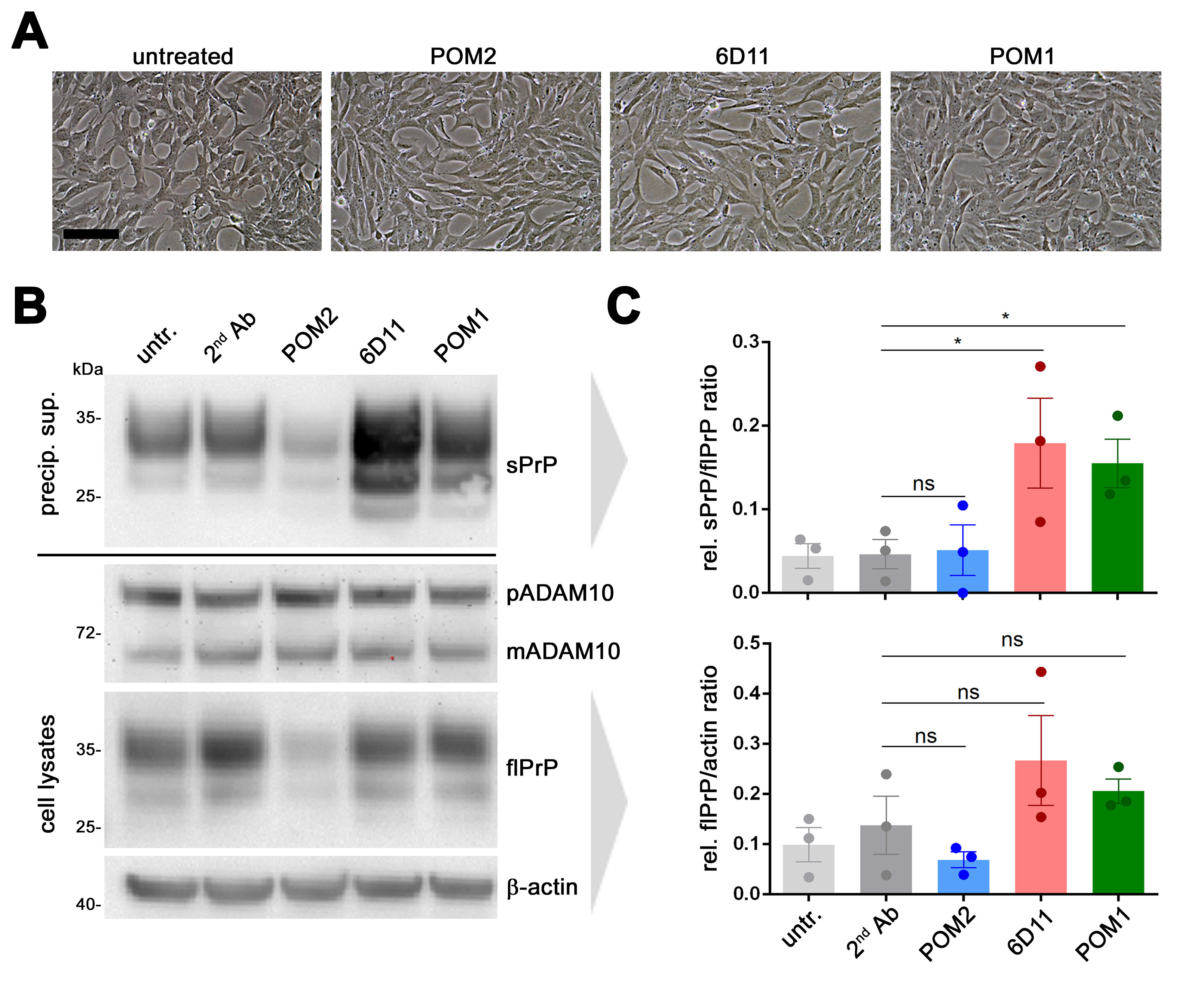

### Suppl Figure 5

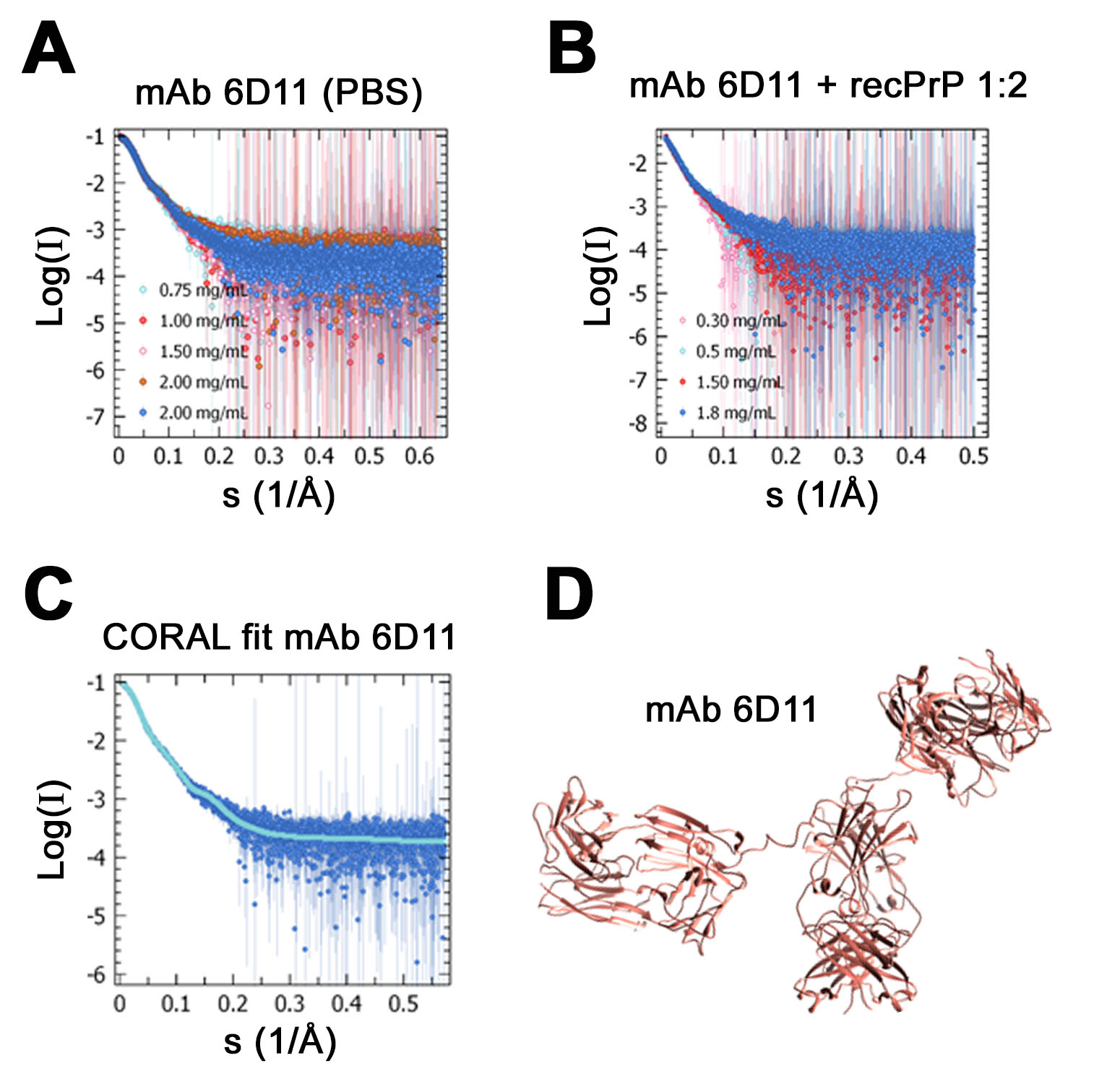

### Suppl Figure 6

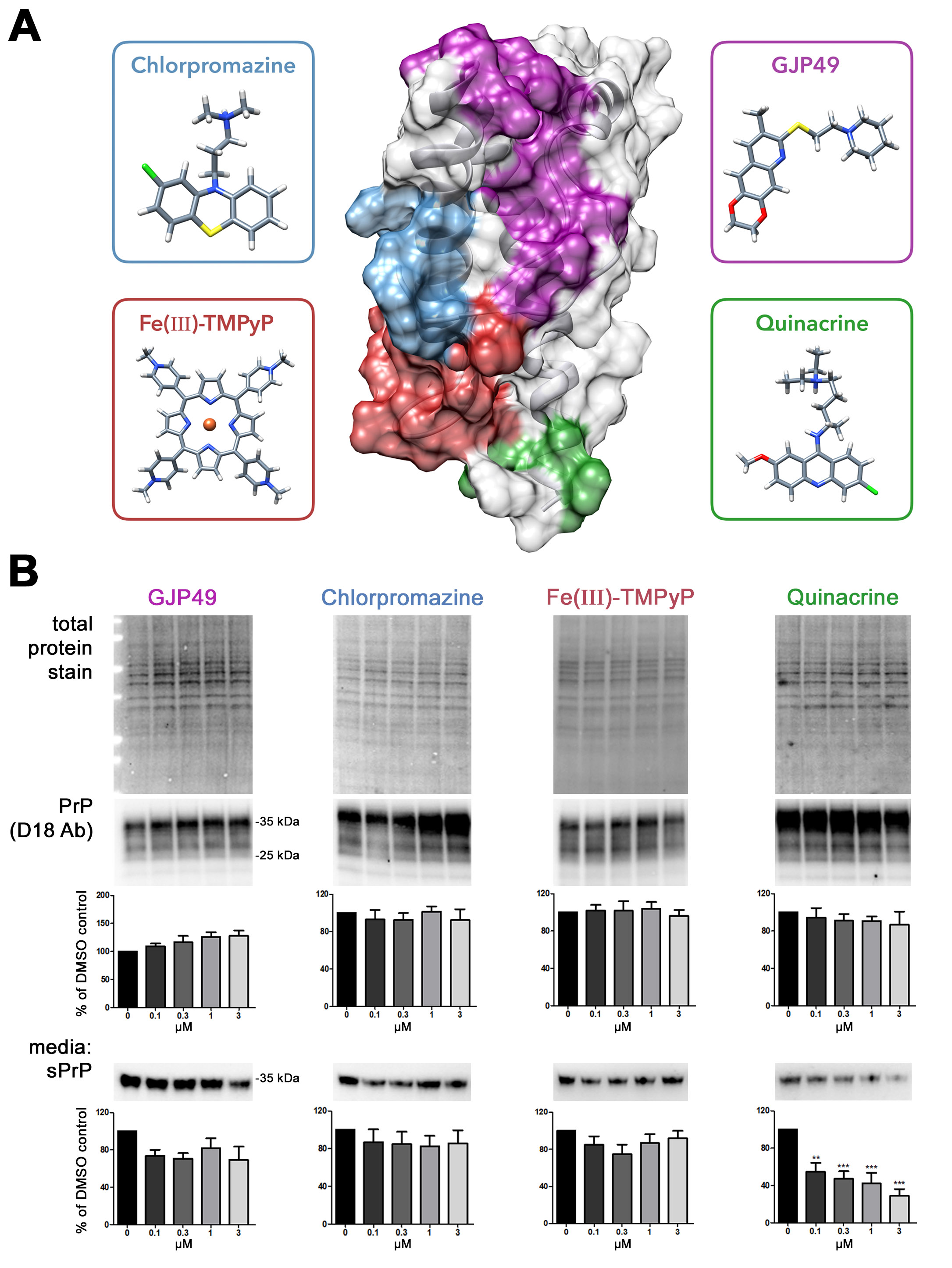

### Suppl Figure 7

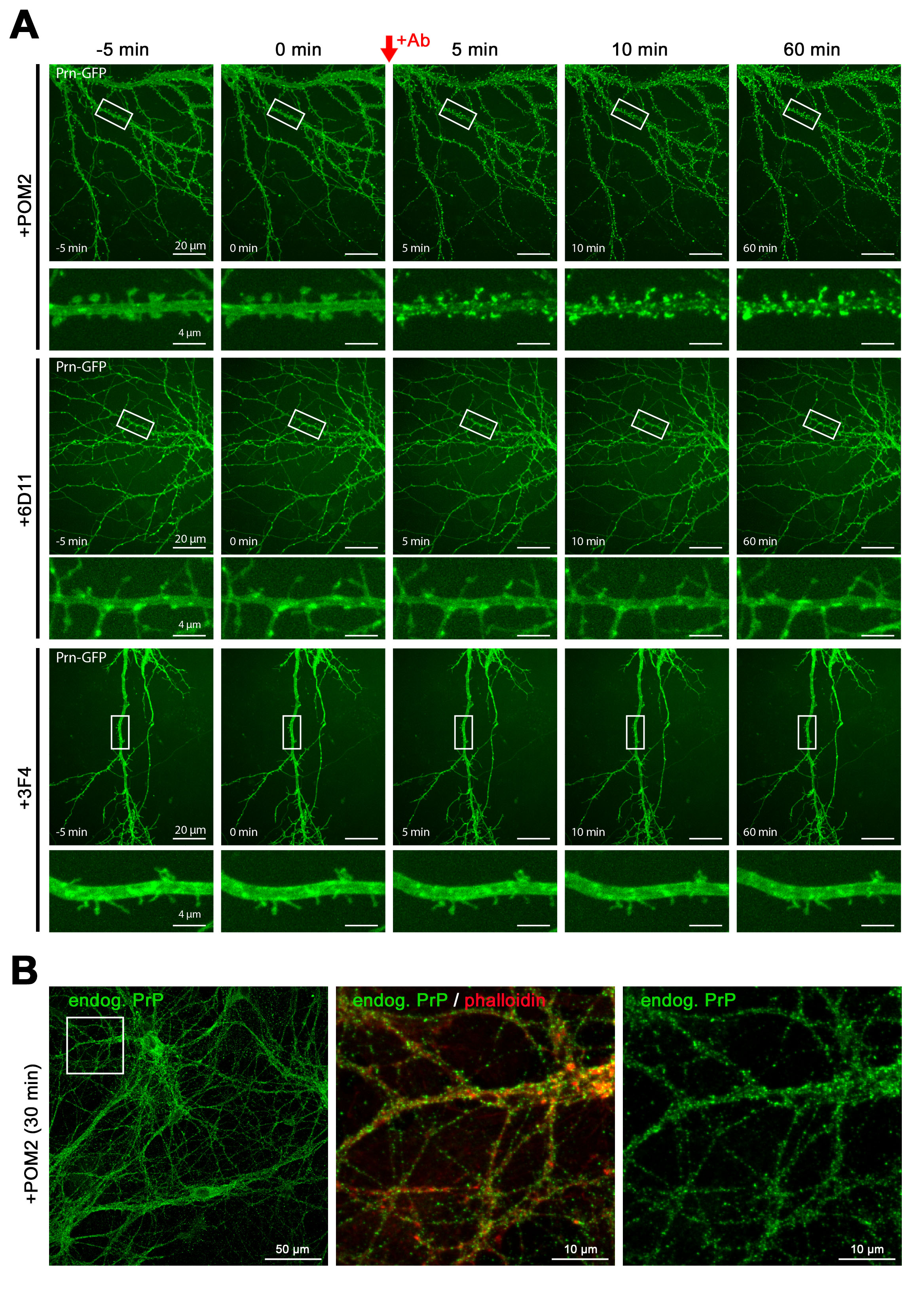

### Suppl Figure 8

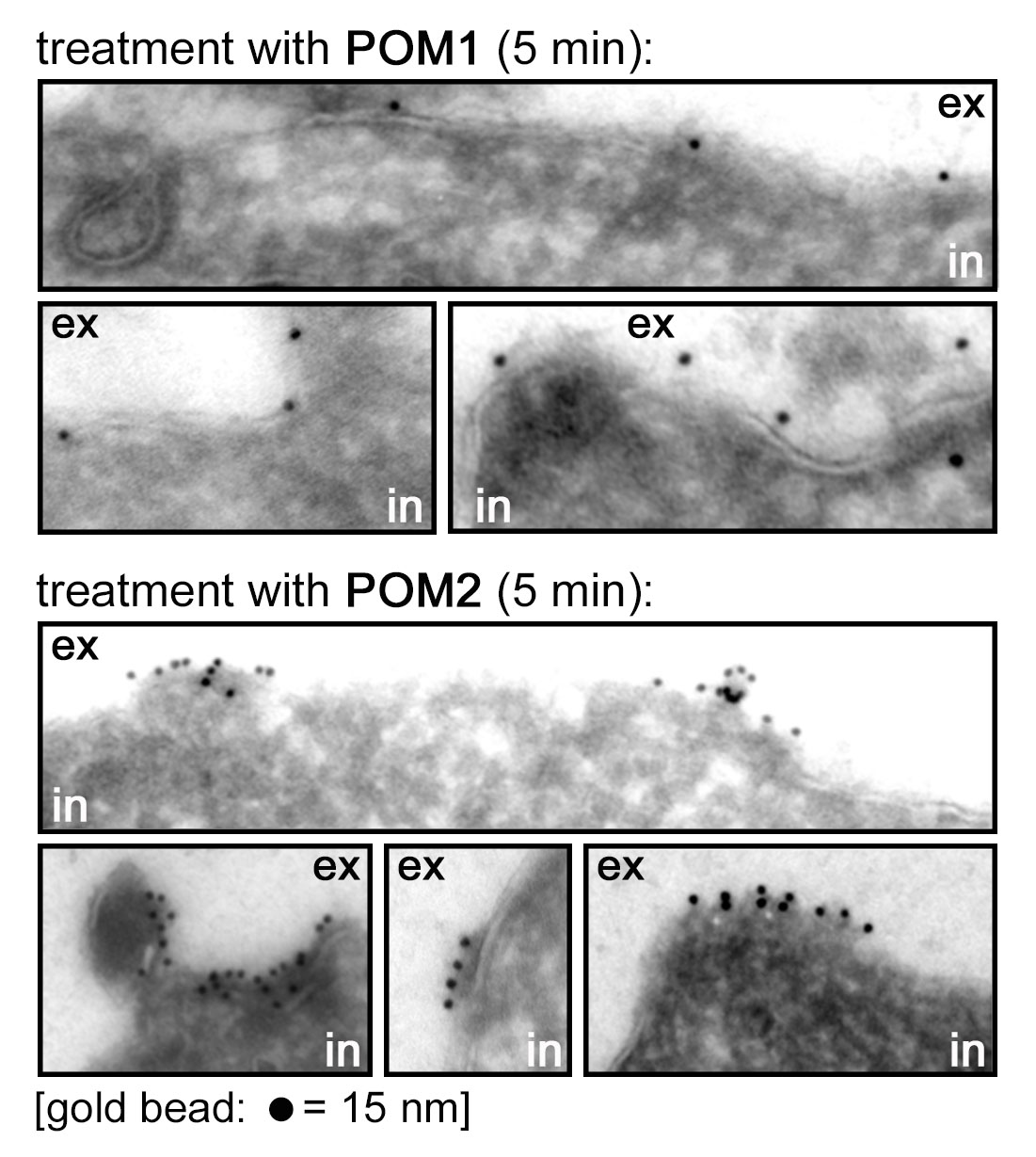
