## Supplementary material for "Ligands binding to the cellular prion protein induce its protective proteolytic release with therapeutic potential in neurodegenerative proteinopathies": Suppl Table 1

**Supplementary Table 1.** Data collection and SAXS-derived parameters

| **Data collection parameters** | | | |
|---|---|---|---|
| Radiation source | Petra III (DESY) | | |
| Beamline | EMBL P12 | | |
| Detector | Pilatus 6M | | |
| Wavelength (nm) | 0.124 | | |
| Sample-detector distance (m) | 3 | | |
| *s* range ($nm^{-1}$) | 0.03 - 7.3 | | |
| Exposure time (s) | 6 (=60 x 0.1 s) or 2 (=20 x 0.1 s) | | |
| Temperature (K) | 293.2 | | |
| **Overall parameters** | **recPrP** | **6D11** | **Complex 2:1** |
| Concentration (mg/mL) | 1.56 | 2.00 | 1.80 |
| $R_g$ from Guinier approximation (nm) | 2.78 ± 0.04 | 5.12 ± 0.03 | 8.13 ± 0.02 |
| $R_g$ from PDDF* (nm) | 2.9 ± 0.1 | 5.23 ± 0.09 | 7.60 ± 0.02 |
| Max.Intramolecular distance $D_{MAX}$ (nm) | 9.9 | 17.6 | 24.8 |
| Porod Volume, $V_P$ ($nm^3$) | 43 | 330 | 954 |
| Molecular weight, I(0) (kDa) | n/a | 120 | 206 |
| Molecular weight from Bayesian estimate (kDa) | 20.8 - 24 | 127 - 151 | 221 - 373 |
| Expected molecular weight (e.g. sequence) (kDa) | 22.9 | 150** | 196 |
| **Software employed** | | | |
| Primary data reduction | SASFLOW | | |
| Data processing | PRIMUS | | |
| *Hybrid (rigid/random loops) and Rigid body* modelling | CORAL, SASpy | | |

*Pair distance distribution function
**Typical IgG molecular mass

### Supplementary information on SAXS measurements and data

SAXS data and modelling for pure **recPrP** accounting for its partial disorder are reported in detail elsewhere (SASBDB accession code: SASDHV9). The overall SAXS-derived parameters of recPrP are presented in Suppl. Table 1, and an ensemble of CORAL models and their best fit to the data are shown in Fig. 4A,C (25% of conformers shown for clarity). The significant variability of the N-terminal parts in these models suggests that recPrP, distal to the membrane anchoring site, samples multiple conformations including very extended ones. The **6D11** antibody is a murine IgG2a. The SAXS profiles of the concentration series displayed in Suppl. Fig. 5 overlap well, thus pointing to the absence of interparticle interactions in the concentration range probed here. In the following, the curve at 2 mg/mL, with the best signal to noise ratio, was employed. The derived overall parameters (Suppl. Table 1, above) are indicative of a typical monomeric IgG. In order to model the conformation of 6D11, the crystal structure of another murine IgG2a [Harris et al. 1997; Ref. 145], PDB accession code 1IGT, was employed. The conformation in the crystal structure does not fit satisfactorily the SAXS data (discrepancy $\chi^2$=2.84, not shown). To obtain representative conformations for the solution state, the model was disconnected in the constituting Fab and Fc domains, and the hinge region of the heavy chains modelled in CORAL as two 20 amino acids long random loops. Ten independent CORAL reconstructions consistently revealed an approximately T-shaped conformation fitting the data well ($\chi^2$=0.84-0.87) (Suppl. Fig. 5), and thus yielding a representative model of the dominant antibody conformer in solution.
The concentration series of the 2:1 recPrP/6D11-antibody complex also shows consistently overlapping curves without concentration-dependent effects (Suppl. Fig. 5B). The complex was modelled against the representative data set collected at 1.8 mg/mL. The overall parameters (Suppl. Table 1) suggest a rather extended assembly and a representative model of a 2:1 complex, with $\chi^2$=0.83, modelled on the basis of these data is shown in Fig. 4D. Intriguingly, to explain the SAXS data, the complex had to be modelled by using very extended conformers for recPrP. While complex formation with such a flexible antigen will almost certainly result in multiple conformations, the solution scattering pattern is dominated by "open" conformations of the complex. This suggests that, in the presence of the 6D11 antibody, PrP (as assessed here for recPrP) tends to adopt predominantly extended conformations.
