## Supplementary material for "Ligands binding to the cellular prion protein induce its protective proteolytic release with therapeutic potential in neurodegenerative proteinopathies": Suppl Table 2

**Supplementary Table 2.** SPT-QD analysis

Median Diffusion Coefficient, D (μm$^2$/s) for data plotted in **Fig. 7B** and **7D**.

Three experiments performed on independent cultures. Total number of QDs analyzed per condition. Kolmogorov-Smirnov test comparing the distribution of diffusion coefficient values.

| | | **POM1 (control)** | **POM1 (1 h)** | **POM19 (control)** | **POM19 (1 h)** | **POM2 (control)** | **POM2 (1 h)** | **POM11 (control)** | **POM11 (1 h)** | **POM3 (control)** | **POM3 (1h)** |
|---|---|---|---|---|---|---|---|---|---|---|---|
| **SYNAPTIC** | **Median D** | 0.1010 | 0.0858 | 0.1008 | 0.0912 | 0.1067 | 0.0555 | 0.0961 | 0.0518 | 0.0956 | 0.0740 |
| | **No. of QDs** | 428 | 591 | 383 | 302 | 213 | 115 | 350 | 288 | 155 | 102 |
| | **KS-test** | ** | | ns | | *** | | *** | | ns | |
| **EXTRA-SYNAPTIC** | | | | | | | | | | | |
| | **Median D** | 0.1594 | 0.1404 | 0.1790 | 0.1505 | 0.1865 | 0.1015 | 0.1560 | 0.0980 | 0.1385 | 0.1421 |
| | **No. of QDs** | 2439 | 2985 | 1757 | 1950 | 915 | 571 | 1192 | 1187 | 770 | 577 |
| | **KS-test** | *** | | *** | | *** | | *** | | *** | |
